# Brain aging differs with cognitive ability regardless of education

**DOI:** 10.1101/2022.02.09.479697

**Authors:** Kristine B. Walhovd, Lars Nyberg, Ulman Lindenberger, Fredrik Magnussen, Inge K. Amlien, Øystein Sørensen, Yunpeng Wang, Athanasia M. Mowinckel, Rogier A. Kievit, Klaus. P. Ebmeier, David Bartrés-Faz, Simone Kühn, Carl-Johan Boraxbekk, Paolo Ghisletta, Kathrine Skak Madsen, Willliam F.C. Baaré, Enikő Zsoldos, Brenda Penninx, Anders M. Fjell

## Abstract

Higher general cognitive ability (GCA) is associated with lower risk of neurodegenerative disorders, but neural mechanisms are unknown. GCA could be associated with more cortical tissue, from young age, i.e. *brain reserve*, or less cortical atrophy in adulthood, i.e. *brain maintenance*. Controlling for education, we investigated the relative association of GCA with reserve and maintenance of cortical volume, -area and -thickness through the adult lifespan, using multiple longitudinal brain imaging cohorts (n = 3327, 7002 MRI scans, baseline age 20-88 years, followed-up up to 11 years). There were widespread positive relationships between GCA and cortical characteristics (level-level associations). In select regions, higher baseline GCA was associated with less atrophy over time (level-change associations). Relationships remained when controlling for polygenic scores for both GCA and education. Our findings suggest that higher GCA is associated with cortical volumes by both brain reserve and -maintenance mechanisms through the adult lifespan.

## Introduction

Does higher intelligence protect against brain atrophy in aging? Numerous findings motivate this question: General cognitive ability (GCA) is positively associated with brain volume and cortical characteristics at various life stages, including young adulthood and older age (1–5). GCA is consistently associated with all-cause mortality and health, with higher GCA related to lower risk of diseases and lifestyle factors known to negatively affect brain health (4). In part, associations are still found after controlling for factors such as educational attainment, suggesting that contemporary GCA in itself is of importance (4). While higher education has been posited as a protective factor against neurodegenerative changes (6, 7), we recently documented in a large-scale study of multiple cohorts that education is not associated with rates of brain atrophy in aging (8). A more promising candidate influence on brain aging may thus be GCA independently of education. Whether GCA level is predictive of longitudinal cortical change has primarily been investigated in older cohorts, and with mixed results (9–11). The relationship of GCA level and cortical changes through the adult lifespan has to our knowledge hitherto not been investigated.

In this context, the lifespan perspective is critical and has implications for understanding functional loss in older age. Several studies indicate that people with higher GCA in young adulthood may be at lower risk of being diagnosed with neurodegenerative disorders in older age (4, 12, 13). Recent findings from large datasets point to a relationship between family history of Alzheimer’s Disease (AD) and cognitive performance level four decades before the typical age of onset of AD(14). However, GCA-AD risk associations have not been consistently observed, and mechanistic factors are poorly understood(15). Possible explanations include both a brain reserve, i.e. “threshold model”(16), as well as a brain maintenance(17) account. The brain reserve model would entail that higher GCA as a trait is related to greater neuroanatomical volumes early in life, young adulthood inclusive, thus delaying the time when people fall below a functional threshold of neural resources in the face of neurodegenerative changes with age. This would happen even if such changes in absolute terms are of similar magnitude across different ability levels, i.e. slopes are parallell (16). The brain maintenance account would on the other hand predict less brain change in adulthood (17) for people of higher GCA, and therefore a smaller risk of cognitive decline and dementia. The brain reserve and maintenance accounts of the relationships between GCA, brain characteristics and clinical risk are not mutually exclusive, but their relative impact through the adult lifespan is unknown. Collectively, the current findings indicate a need to understand whether there is a relationship between GCA as a trait and brain changes, independently of education, over the adult lifespan.

We tested whether GCA predicted brain aging as indexed by cortical volume, area and thickness change measured longitudinally in 7084 MRI scans from several European cohorts covering the adult lifespan in the Lifebrain consortium(18) and the UK Biobank (UKB) (19, 20) (n = 3327, age range 20-88 years at baseline, maximum scan interval of 11 years, see Online Methods for details). To disentangle possible environmental and genetic influences on the relationship between GCA and brain aging, we controlled for educational attainment in the main analyses, and in a second step for polygenic scores (PGSs) for education and GCA (21, 22).

Established PGSs are only moderately predictive of GCA (21), but in view of evidence that the polygenic signal clusters in genes involved in nervous system development (21), we did expect such scores to explain part of the intercept effect, with no or weaker effects on brain aging. We expected any effects of GCA on cortical changes to apply to all ages, but in view of recent findings of greater relationships between brain and cognitive function in older than younger individuals(3), we also tested the age interaction. Based on previous findings, including from broader cross-sectional Lifebrain cohorts (23), and mixed results from smaller longitudinal older cohorts (9–11), we hypothesized that GCA would be positively related to anatomically widely distributed cortical characteristics through the adult lifespan (intercept effect), but that associations with differences in cortical aging trajectories (slope effects) may be observed to a lesser extent. We expected effects of GCA to be at least partially independent of education(8), both for intercept and slope associations.

## Results

The main models of associations of GCA with cortical characteristics, and their change, were run separately for samples within the Lifebrain consortium (n = 1129, 2606 scans) (18) and the UK Biobank (UKB, n = 2198, 4396 scans)(19, 20). In all main models, sex, baseline age, scanner, time (interval from baseline) and education were entered as covariates. In modeling the effects of GCA on cortical characteristics *(level-level analyses)*, GCA was entered as the predictor (explanatory variable), whereas in modeling the effects of GCA on brain aging *(level-change analyses)*, the interaction term of GCA x time was entered as the predictor, and education x time was entered as an additional covariate along with GCA and education. Since brain aging (i.e. change) was of chief interest, we did not include intracranial volume (ICV), which is stable, in the main analyses. Results from models including ICV, as well as models without education, as covariates, can be found in the Supplemental Information (SI). Additional analyses included the interaction term baseline age x time as a covariate, and in one set of analyses we entered the interaction term baseline age x time x GCA as predictor (with relevant two-way interaction terms as covariates), to test if effects differ reliably across the lifespan.

### GCA level – brain level analyses

P-value maps for the relationship of GCA and cortical characteristics controlled for education, are shown in Figure 1. For cortical volume and area, there were widespread positive effects of GCA bilaterally across the cortical mantle in UKB. Largely overlapping, but more restricted effects were seen in the Lifebrain sample. Uniformly across samples, higher GCA was associated with greater cortical volume and area. For cortical thickness, only minor positive effects were seen in UKB, in proximity of the left central sulcus. Analyses not controlling for education are shown in Supplemental Figure 1, with similar effects for cortical volume and area, but with a negative association for cortical thickness in the right anterior cingulate area in the UKB. The introduction of the interaction term of age x time as a covariate in the main analyses shown in Figure 1, did not influence results strongly (see Supplemental Figure 2), indicating that the observed relationships may be relatively uniform throughout the adult lifespan. When adding ICV as a covariate, the intercept effects in the analysis shown in Figure 1 became non-significant in UKB, pointing to these being broad effects grounded in greater neuroanatomical structures in general, rather than being region-specific. However, the more restricted regional intercept associations of GCA and cortical volume and area largely remained in the Lifebrain samples also when adding ICV as an additional covariate (Supplemental Figure 3).

**Figure 1.**
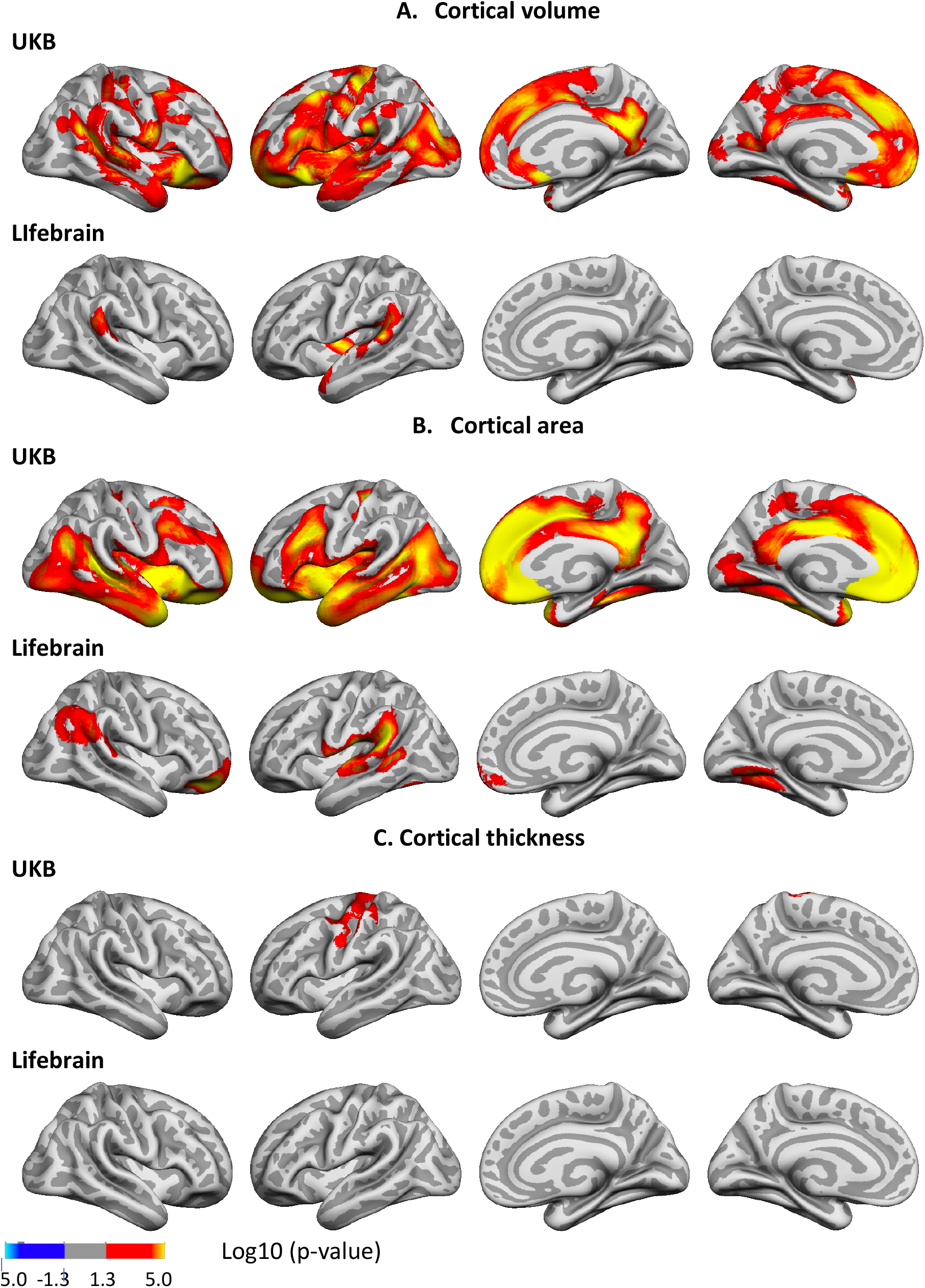
P-value maps of the associations of general cognitive ability (GCA) and cortical characteristics, controlled for education. P-value maps of the relationships between GCA at baseline and cortical characteristics are shown, when age at baseline, sex, time (since first scan) and education are controlled for (p <.01, corrected using a cluster-forming p-value threshold of p < .01). Relationships are shown, from left to right for each panel: right and left lateral view, right and left medial view, for UKB and Lifebrain samples for A. Cortical volume, B. Cortical area, and C. Cortical thickness.

### Gca level- brain change analyses

Having confirmed the expected positive relationships between GCA and cortical volume and area controlled for education in terms of an intercept effect, we investigated the question of slope effects was next. Associations of GCA level at baseline and change in cortical characteristics, controlled for education, are shown in Figure 2. Higher baseline GCA was associated with less regional cortical volume reduction and thickness reduction over time in both UKB and Lifebrain. As expected, effects were more spatially limited than those seen for intercept models. For volume, effects did not overlap between samples. In UKB, left medial parietal and frontal effects were identified, whereas in Lifebrain, left lateral occipital and right medial occipitotemporal effects were observed, and right medial and lateral frontal, and medial occipitotemporal associations were observed. All volume change associations were positive (i.e. higher GCA was associated with less volume reduction). No associations with area change were observed in UKB, whereas a small region in the right parietal cortex showed a negative association, and a small region in the lateral frontal cortex showed a negative association in Lifebrain. For cortical thickness, somewhat broader and bilateral positive effects were seen in both samples, partially overlapping in lateral temporal and posterior areas as well as medial posterior and frontal areas of the left hemisphere. Taken together, this means that the observed positive associations of GCA with volume change primarily reflect less cortical thinning with higher GCA. It should be noted, that at a more lenient threshold of correction for multiple comparisons (corrected at p <.05, cluster-forming threshold of p<.05), associations of GCA with cortical thickness change were overlapping to a greater extent. Partially overlapping left medial posterior and anterior associations with volume changes, and bilateral associations with thickness changes, partially overlapping medially, were at this more lenient significance threshold observed both in UKB and Lifebrain (Supplemental Figure 4).

**Figure 2.**
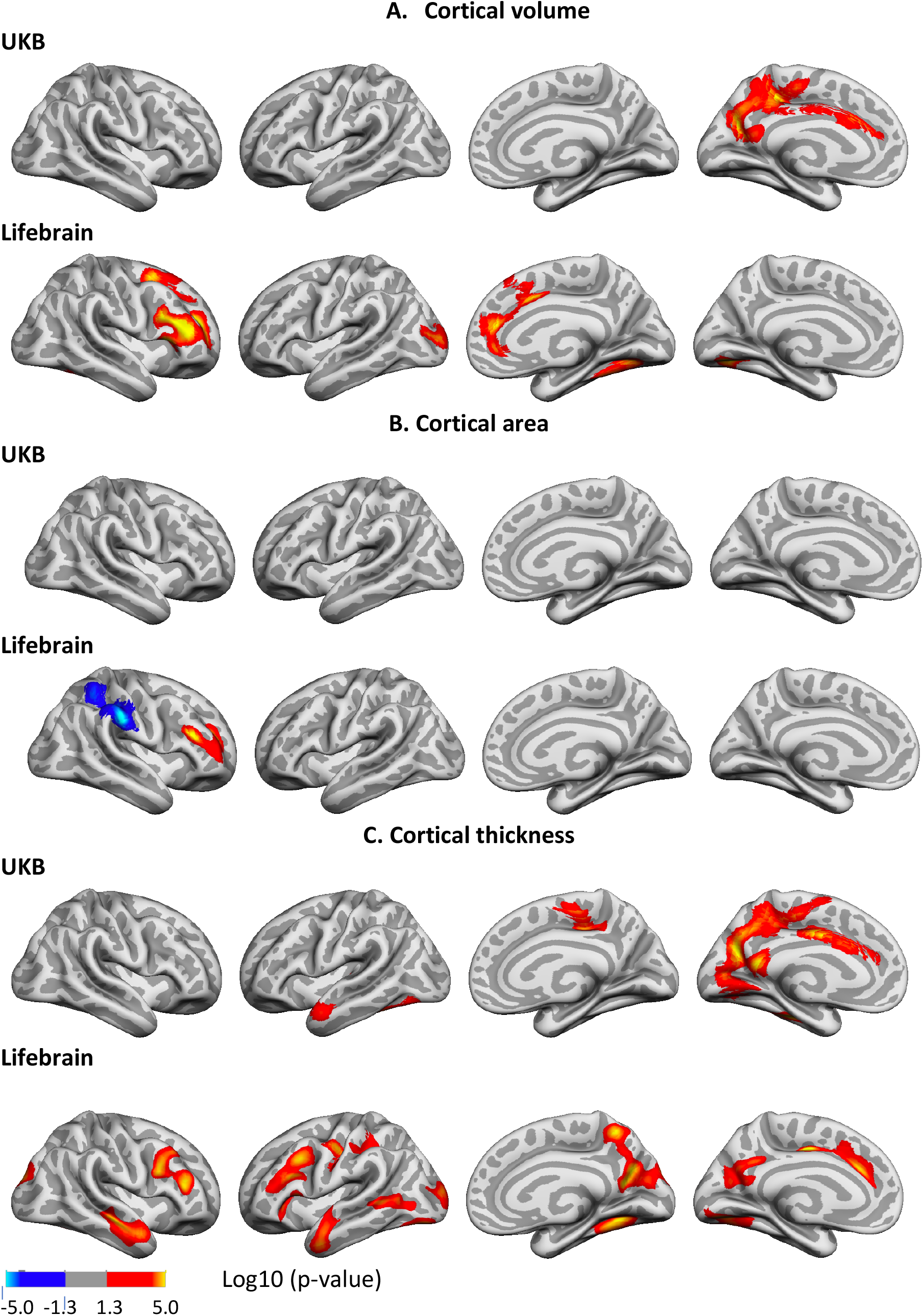
P-Value maps of the associations of general cognitive ability (GCA) with change in cortical characteristics, controlled for the effect of education over time. The association is shown for the interaction of GCA at baseline and time (interval since baseline scan), when age, sex, time, GCA, education, and the interaction of education and time, are controlled for (p <.01, corrected using a cluster-forming p-value threshold of p<.01). Significant regions are shown, from left to right for each panel: right and left lateral view, right and left medial view, for UKB and Lifebrain samples: A. Cortical volume, B. Cortical area, C. Cortical thickness.

The associations of GCA with cortical change were essentially unaffected by adding ICV as a covariate (Supplemental Figure 5).

In order to characterize consistency of effects across samples (UKB and Lifebrain), we plotted the generalized additive mixed model (GAMM) for the different GCA quintiles, from lowest to highest (Figure 3), depicting change trajectories for average cortical volume and thickness within the regions showing significant GCA x time associations. Across samples, subgroups with higher GCA started with higher volume *and* had less volume loss over time. For instance, on average, people with maximum cognitive score in UKB are expected start out with a regional average cortical volume of 1.72 mm^3^ that would be maintained for the next three years, whereas those with the lowest GCA would on average start out with 1.64 mm^3^ and decrease to 1.61 mm^3^ over the next three years. Thus, the greatest GCA-associated differences in cortical volume are found in the intercepts (level), whereas differences in slope (change) are smaller in the follow-up period. For cortical thickness, the change trajectories were also very consistently ordered, but those with higher GCA did not uniformly have thicker cortex at first timepoint in these areas. Rather, differential rates of cortical thinning over time were critical in creating cortical thickness differences in these regions in aging. This was evident in both samples, but especially pronounced in UKB.

**Figure 3.**
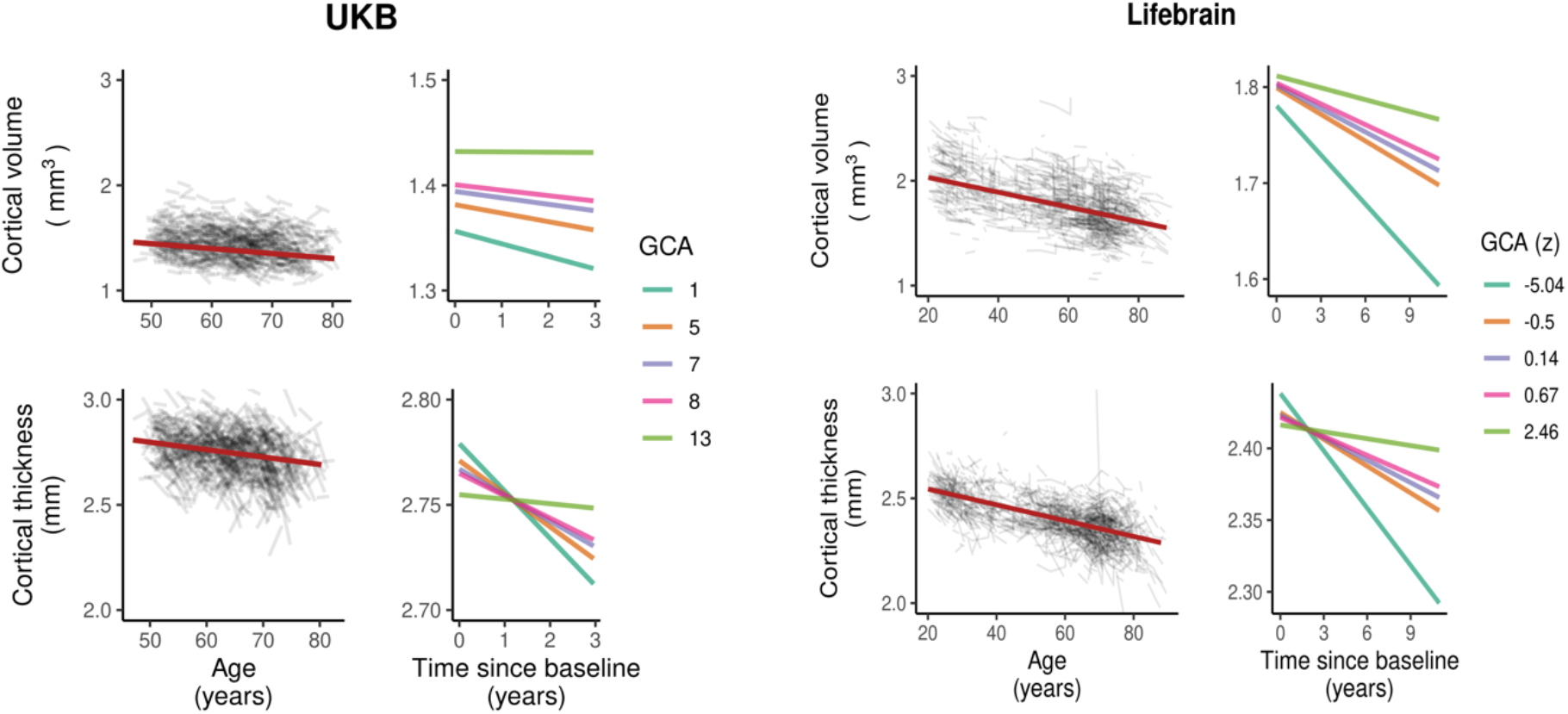
Cortical change trajectories according to general cognitive ability (GCA). Trajectories are shown per quintile of GCA, for mean cortical volume and thickness change in the analysis model and regional significant sites of associations shown in Figure 2. For UKB, the quintiles refer to actual scores from min to max (0-13) on the test, whereas for Lifebrain, the quintiles refer to z-scores (where mean is zero) min to max for the samples.

### Influence of polygenic scores (PGSs) for GCA and education on the level-level and level-change associations

Next, we investigated whether effects were maintained when covarying for established PGSs for GCA and education in the UKB(21, 22). Fifty-two participants in the main models were excluded due to missing genetic data. In these analyses, we regressed out the first ten genetic ancestry factors (GAFs) from the GCA variable prior to analysis. The intercept associations of GCA and cortical characteristics that were observed in the main model (Figure 1) largely remained when controlling for the PGSs, but the extent of the significant regions were somewhat reduced for cortical volume and area (Supplemental Figure 6). The associations of GCA and cortical change largely remained and were only slightly attenuated when controlling for PGSs for GCA and education (Figure 4).

**Figure 4.**
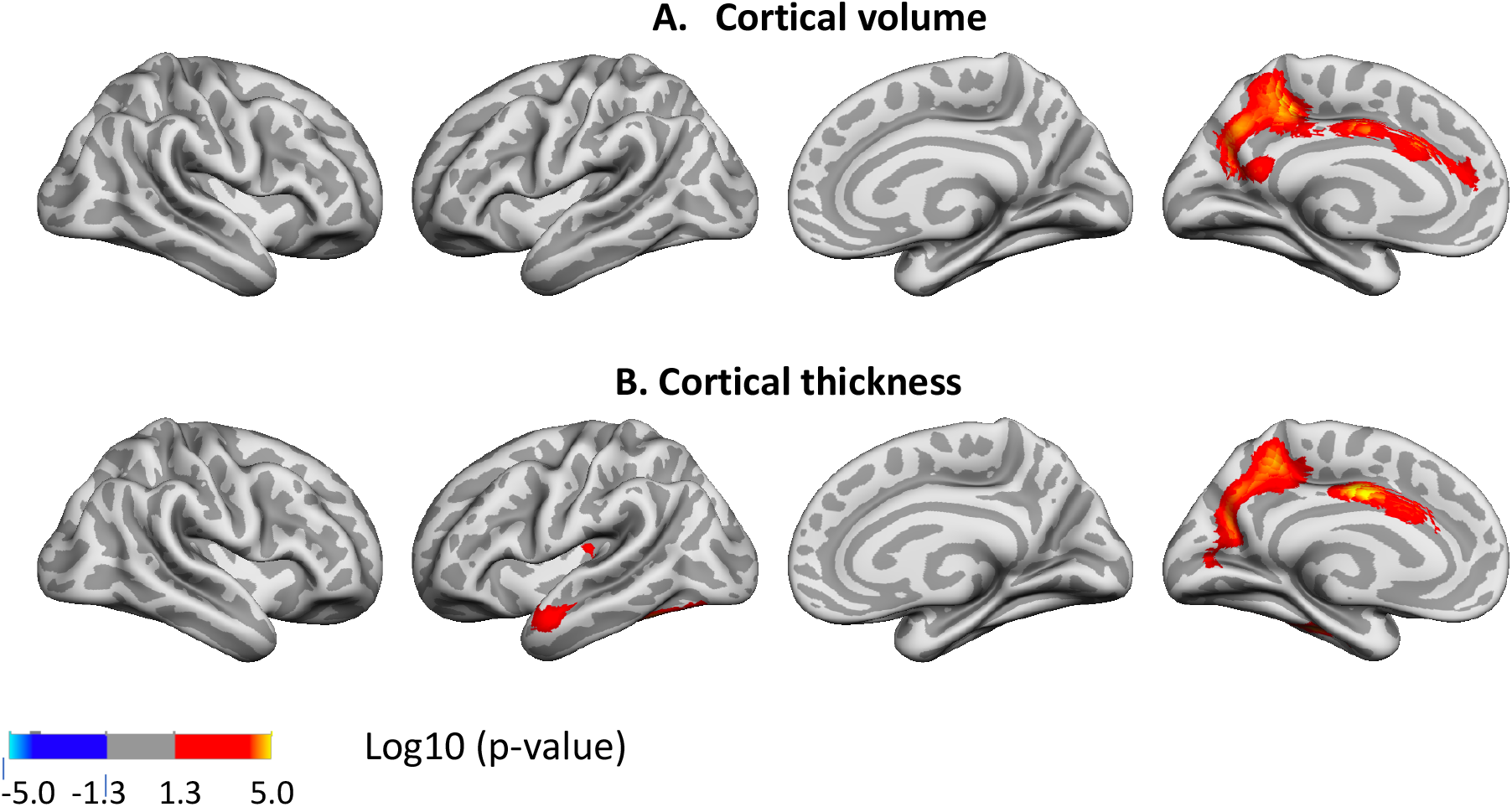
P-value maps of associations of general cognitive ability (GCA) and change in cortical characteristics in the UKB, controlled for the effect of education and polygenic scores (PGSs) for education and GCA over time. The significant regions are shown for the interaction of GCA at baseline (with genetic ancestry factors regressed out) and time (interval from baseline scan), when age, sex, time, GCA, education, and the interactions of education by time, and PGSs by time, are controlled for (p <.01, corrected using a cluster-forming threshold of p<.01). Regions are shown, from left to right for each panel: right and left lateral view, right and left medial view, for UKB and Lifebrain samples: A. Cortical volume, B. Cortical thickness. No significant regions were seen in either sample for cortical area.

### Influence of age on the level-change associations

We next tested the three-way interaction baseline GCA x baseline age x time, to see whether the level-change associations differed reliably across the lifespan. In UKB (age range 47-80 years), no such interaction effects were observed, but in the more age-heterogeneous Lifebrain cohort (age range 20-88 years), significant interaction effects were seen for change in small regions of the left hemisphere laterally for volume, and for slightly more extended bilateral lateral regions, as well as for a left posterior fusiform region for cortical thickness (see Figure 5).

**Figure 5.**
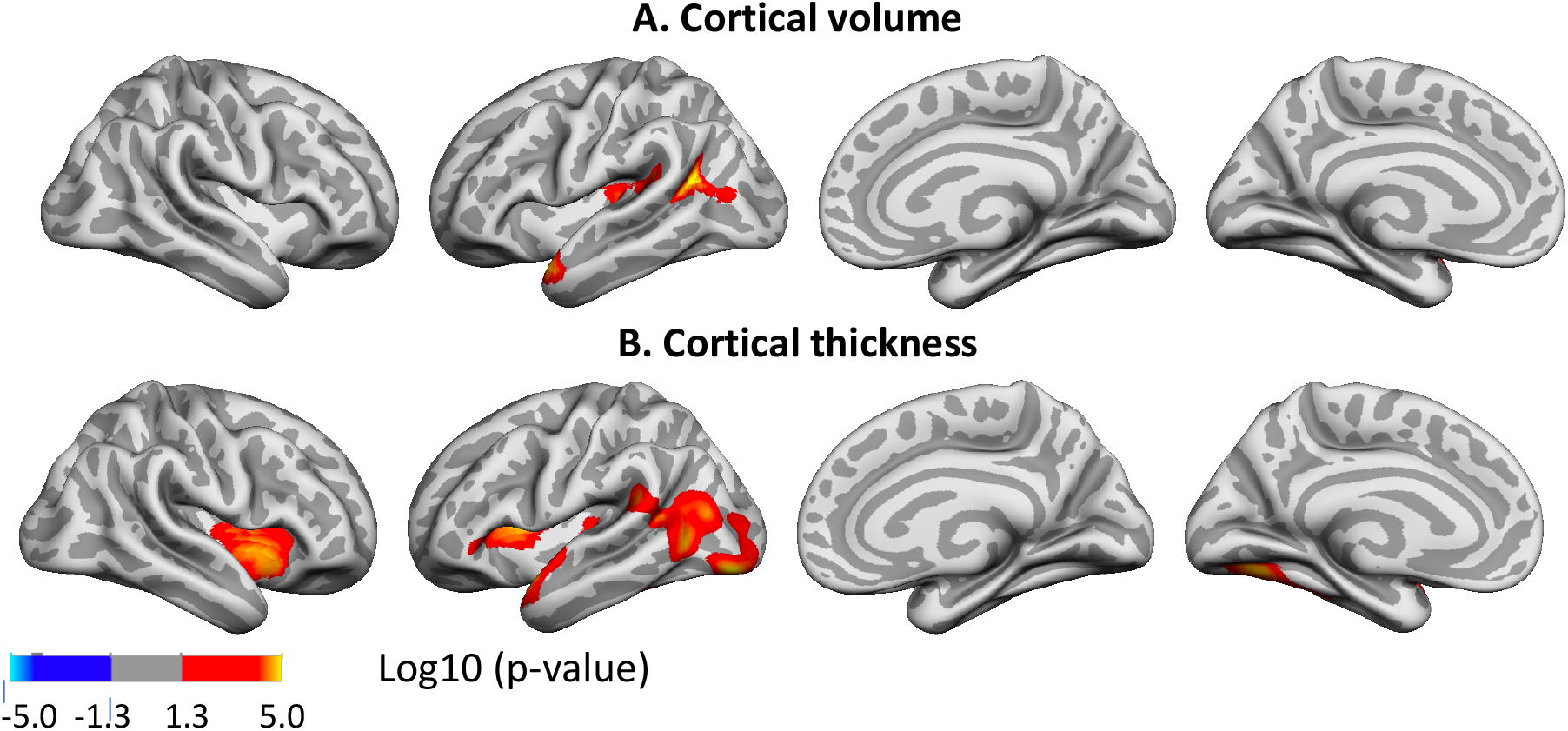
P-value maps for interactions of level general cognitive ability (GCA), by baseline age by time on cortical volume and thickness in the Lifebrain cohort. The significant regions are shown for the interaction of GCA at baseline, by age at baseline by time (interval since baseline scan), when age, sex, time, GCA, education, and the interactions education by time, GCA by time, and baseline age by time, are controlled for (p <.01, corrected using a cluster-forming threshold of p<.01). Effects are shown, from left to right: right and left lateral view, right and left medial view, for A. Cortical volume, and B. Cortical thickness, in the Lifebrain sample. No significant regions were seen for cortical area in Lifebrain, and no significant regions were seen in UKB.

In the Lifebrain cohort, the positive three-way interactions of baseline GCA x baseline age x time indicates that higher level of GCA is associated with less atrophy at distinct ages. To visualize these interaction effects, we divided the cohort according to whether participants were above or below age 60 years. This division point was chosen in view of it being an approximate age at which select cognitive and regional cortical volume and thickness changes have been reported to accelerate in longitudinal studies(24, 25). In addition, the subsamples were approximately equally powered (1293 scans for the younger versus 1313 scans for the older half). In order to explore how GCA level related to cortical thickness change over time across the two age groups, we plotted the expected cortical change trajectories as a function of GCA, with each sample divided into quintiles, from lowest to highest GCA. The plots are shown in Figure 6. While GCA level was not or weakly inversely related to atrophy in these regions in the younger group, the expected trajectories for the older group were relatively consistently ordered so that persons with a higher GCA level had less decline of cortical thickness. Figure 6, lower panel shows that that quintile differences are more pronounced in the older group, suggesting the latter half of the lifespan is driving the interaction. As one outlier in the older group was noted as having a high cortical thickness value for age at the first timepoint in the region of interest, we carefully checked this segmentation, but found no sign of flawed segmentation, and thus decided to keep this person in analyses.

**Figure 6.**
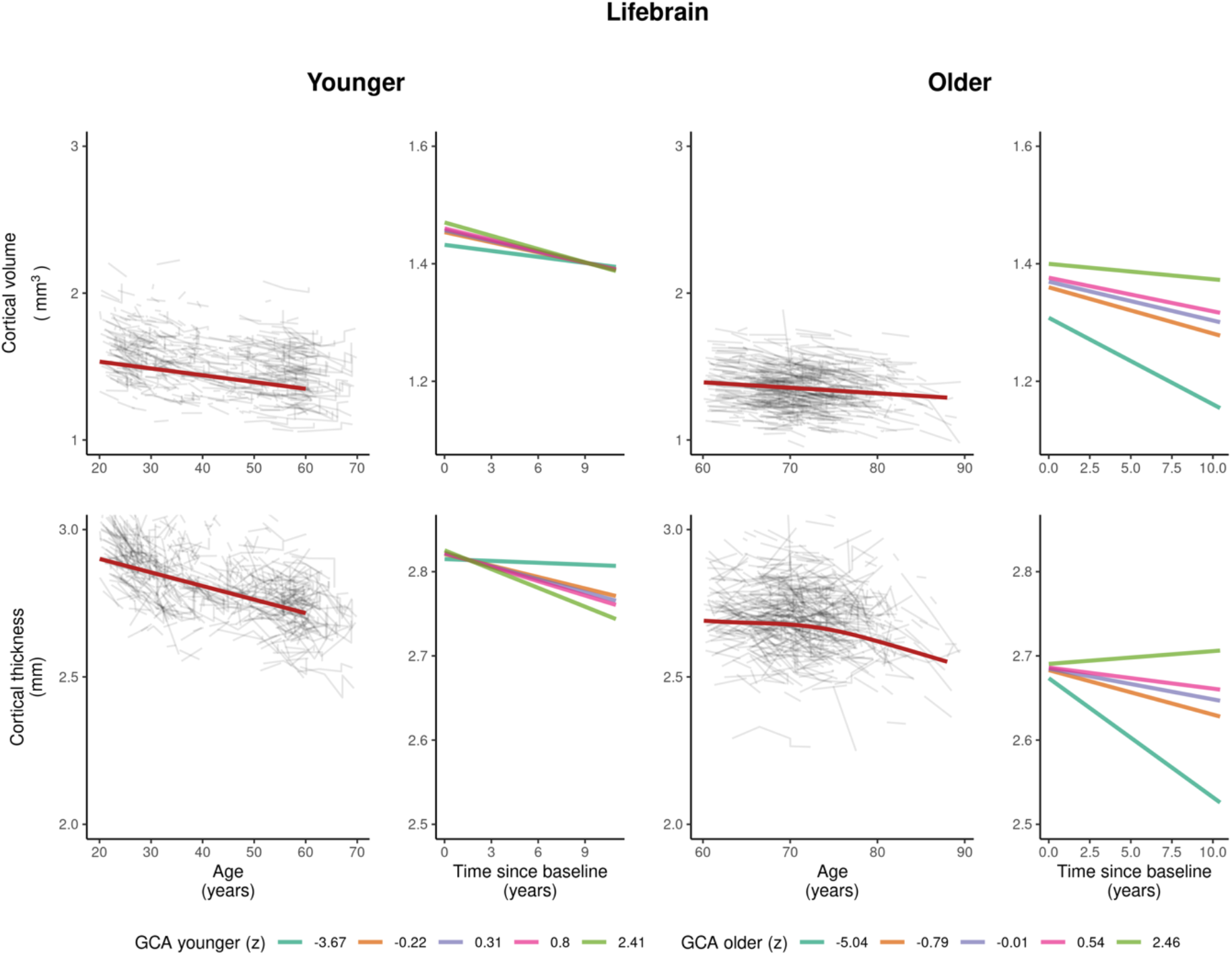
Cortical volume and thickness change trajectories according to general cognitive ability (GCA) for young and older adults in the Lifebrain cohort. Expected trajectories are shown for young (< 60 years at baseline) and older (> 60 years at baseline) adults in the Lifebrain cohort per quintile of GCA, for mean cortical thickness in the analysis model and regional significant sites shown in Figure 5. The quintiles refer to z-scores min to max for the sample.

## Discussion

The current study provides novel findings on GCA not only as a marker of brain characteristics, but also of brain changes in healthy aging. The finding that higher GCA level is associated with larger neuroanatomical structures to begin with, i.e. greater brain reserve, confirms findings in previous studies of various age groups(1–4, 23, 26). While level of GCA has been associated with cortical change in some older groups(10, 11), but not others(9), the current demonstration of an association of GCA levels, controlled for education, on cortical volume and -thickness declines through the lifespan in multiple cohorts across a long follow-up time and multiple follow-ups, constitutes a novel finding. Also, the finding of an age -interaction with pronounced effects of GCA on cortical thinning and volume changes only in older ages in select regions in the more age-heterogeneous Lifebrain cohort, is novel.

The association of GCA level and cortical change appears relatively moderate. This may explain why such associations have not previously been consistently found. The “effect” of GCA on cortical change must be viewed in relation to the intercept effects, which, as shown here, constitute a major source of GCA-related cortical volume variation through the lifespan: Those with higher GCA have greater cortical area to begin with, yielding higher cortical volumes in young adulthood. We have previously found that cortical area seems in part determined neuro-developmentally early on, is associated with GCA, and shows parallel trajectories for higher and lower GCA groups.(1) As there is a relatively minor age change in area, compared to thickness(25), slope effects on cortical volume are chiefly caused by moderately different rates of cortical thinning for people of differential cognitive ability. Differences in cortical thinning are thus key to the maintenance effects of GCA, whereas early differences in cortical area drive the intercept effect. Through the adult lifespan, both will affect cortical volume.

It is of interest that these GCA-brain change associations were found when education was controlled for, suggesting that the contemporary GCA level may not only be related to brain reserve to begin with, but also brain maintenance. This is evident from the – across samples –consistently steeper slopes of regional cortical decline with lower GCA (as shown in Figure 3). With our recent findings on the variable nature of education-brain-cognition relationships, as well as education not being associated with atrophy rates in aging(23, 27), this points to the component of GCA not being associated with education variance as a more promising candidate for predictive or potentially protective effects on brain aging. There is evidence that education may serve to increase GCA(28, 29). However, while GCA level may be impacted, slope, i.e. cognitive decline, is likely not(29, 30). There is also evidence to suggest that education, without mediation through adult socioeconomic position, cannot be considered a modifiable risk factor for dementia(31). While one would then think the underlying mechanism in the observed GCA-brain change relationships may be genetic, known genetic factors only partially explained relationships, as effects remained after controlling for PGSs for general cognitive ability and education. However, the PGSs are known to be only moderately predictive of GCA(21), and genetic pleiotropic effects on GCA and cortical characteristics and their change may still likely be part of the underlying mechanism. While it has been suggested that GCA may associate with differences in epigenetic age acceleration, it was recently reported that such epigenetic markers did not show associations with longitudinal phenotypic health change(32). While it is possible that individual differences in epigenetic age acceleration in older age could be caused by e.g. behaviors associated with intelligence differences over the life course, differences in epigenetic markers and GCA could also both be the result of a shared genetic architecture or some early, including in-utero, environmental event(32, 33).

While controlling for the interaction of baseline age and time had little effect in the level-level analyses, indicating at least partially stable associations through the adult lifespan, a significant three-way interaction of baseline GCA by baseline age by time on regional cortical thickness changes was found in the Lifebrain cohort. These effects indicated that higher level of GCA is more associated with less atrophy at older ages. However, as the significance of these regional interaction effects also seemed to rest in part on unexpected, albeit weakly, inverse direction of smaller effects in younger(25) age, and no such effects were seen in UKB, we consider them tentative until replicated. The absence of age interaction in the UKB could be due to the higher baseline age of the sample, making it unsuitable to study adult lifespan interactions. Moreover, greater power would be desired to study three-way interactions of possibly smaller effect size.

Finally, change-change relationships between GCA and cortical characteristics could not readily be addressed in the present samples with similar models, due to variability in availability of comparable test data across timepoints. In a lifespan perspective, we know that such relationships do exist, in that both brain and cognition increase in development and decline in aging(17, 24, 25, 34). However, to what extent individual differences in GCA *change* are related to individual differences in cortical trajectories in the present samples, is beyond the scope of this study.

In conclusion, the present study shows that with higher GCA, primarily brain reserve, but also brain maintenance yield higher cortical volumes through the adult lifespan. These effects were seen when controlling for effects of education. As there is otherwise scarce evidence so far that human behavioral traits are associated with differential brain aging trajectories, this is of great interest to investigate further. While controlling for known polygenetic markers for GCA and education did not substantially diminish the effects, the underlying mechanisms may still be related to genetic pleiotropy. However, this leaves open the possibility that factors associated with increased GCA other than education, and possibly genes, could serve to diminish cortical atrophy in aging. Such factors affecting normal individual differences in GCA are not known with certainty, but as childhood GCA is highly predictive of GCA in aging(35), they likely work at developmental, rather than adult and senescent stages.

## Materials and methods

The UK Biobank (UKB)(20) and the Lifebrain samples are described in Table 1. The samples from the European Lifebrain (LB) project (http://www.lifebrain.uio.no/)(18) included participants from major European brain studies: the Berlin Study of Aging II (BASE II) (36), the BETULA project(24), the Cambridge Centre for Ageing and Neuroscience study (Cam-CAN)(37), Center for Lifebrain Changes in Brain and Cognition longitudinal studies (LCBC)(1), and the University of Barcelona brain studies (UB)(38–40)(6–8).

**Table 1.** Overview of sample characteristics of included cohorts. Age, education, and scans intervals (since baseline) are given in years. GCA for Lifebrain is standardized per sample for first timepoint. GCA is given in z-scores for Lifebrain (see Supplemental Methods), and test-scores for UKB.

| **Sample** | **Lifebrain** |  |  | **UKB** |  |  |
| --- | --- | --- | --- | --- | --- | --- |
| **N** | 1129 |  |  | 2198 |  |  |
| **participants** |  |  |  |  |  |  |
| **N Females** | 576 |  |  | 1136 |  |  |
| **N scans** | 2606 |  |  | 4396 |  |  |
|  | **M** | **SD** | **Range** | **M** | **SD** | **Range** |
| **Baseline age** | 55.5 | 18.4 | 20.0-88.0 | 20.0 | 7.1 | 47.0-80.3 |
| **Scan interval** | 3.7 | 2.7 | 0.2-11.0 | 2.3 | 0.1 | 2.0-3.0 |
| **Education** | 14.9 | 3.1 | 2.0-20.0 | 14.0 | 2.3 | 10.0-16.0 |
| **GCA** | 0.0 | 1.0 | -5.0-2.5 | 6.8 | 2.0 | 1.0-13.0 |

GCA was measured by partially different tests in the different cohorts. National versions of a series of batteries and tests were used, see SM for details. These included the UKB Fluid Intelligence test(41), tests from the Wechsler batteries(42–44) combined with the National Adult Reading Test (NART)(45), the Cattell Culture Fair Test(46) combined with the Spot The Word task(47), as well as local batteries, for which procedures are described in SM and elsewhere(48, 49). It is clearly a limitation that content and reliability of the GCA measures may vary, but there is reason to assume that the measures index partally similar abilities. For instance, the UKB fluid intelligence measure has been shown to have moderate to high reliability, and correlated >.50 with a measure of GCA created using 11 reference tests, including NART and Wechsler measures (50). See SM for further details.

MRIs were processed using FreeSurfer, version 7.1 for Lifebrain, and version 6.0 for UKB (https://surfer.nmr.mgh.harvard.edu/)(51–54). We ran vertex-wise analyses to assess regional variation in the relationships between cortical structure and the measures of interest, i.e. GCA and the interaction of GCA ×time. Cortical surfaces were reconstructed from the same T1-weighted anatomical MRIs, yielding maps of cortical area, thickness and volume. Surfaces were smoothed with a Gaussian kernel of 15 mm full-width at half-maximum. Linear mixed models were run (using FreeSurfers ST-LME package https://surfer.nmr.mgh.harvard.edu/fswiki/LinearMixedEffectsModels), for each of the samples separately, with GCA, and then additionally with the interaction term of GCA and time in turn as predictors, and, sex, baseline age, scanner, time (interval since baseline scan) and education were entered as covariates unless otherwise noted. The results were thresholded at p <.01 (corrected).

All participants gave informed consent, subprojects were approved by the relevant ethical review boards, and the Lifebrain project was approved by Regional Committees for Medical Research Ethics–South East Norway. Screening criteria were not identical across studies, but participants were recruited to be cognitively healthy and did not suffer from neurological conditions known to affect brain function, such as dementia. All samples consisted of community-dwelling participants, some were convenience samples, whereas others were contacted on the basis of population registry information. Further details on samples, GCA measures, MRI acquisition and processing and statistical analyses, are presented in SM. The Lifebrain data supporting the results of the current study are available from the PI of each sub-study on request (see SM), given approvals. UK Biobank data requests can be submitted to http://www.ukbiobank.ac.uk. Computer code used for the analyses is available on github: https://github.com/Lifebrain/p032-gca-brain-change

## Acknowledgements and funding sources

The Lifebrain project is funded by the EU Horizon 2020 Grant agreement number 732592 (Lifebrain). In addition, the different sub-studies are supported by different sources: LCBC: The European Research Council under grant agreements 283634, 725025 (to A.M.F.) and 313440 (to K.B.W.), as well as the Norwegian Research Council (to A.M.F., K.B.W.), The National Association for Public Health’s dementia research program, Norway (to A.M.F). Betula: a scholar grant from the Knut and Alice Wallenberg (KAW) foundation to L.N. Barcelona: Partially supported by an ICREA Academia 2019 grant award; by the California Walnut Commission, Sacramento, California. BASE-II has been supported by the German Federal Ministry of Education and Research under grant numbers 16SV5537/ 16SV5837/ 16SV5538/ 16SV5536K /01UW0808/ 01UW0706/ 01GL1716A/ 01GL1716B, the European Research Council under grant agreement 677804 (to S.K.). The Cambridge Centre for Ageing and Neuroscience (Cam-CAN) was supported by a programme grant from the UK Biotechnology and Biological Sciences Research Council (grant number BB/H008217/1) and by continued intramural funding from the UK Medical Research Council to the Cognition & Brain Sciences Unit in Cambridge. Part of the research was conducted using the UK Biobank resource under application number 32048.

## Author Contributions

KBW drafted the manuscript, after critical discussions of the research questions tested with LN and AMF. IKA, FM, ØS, YW and AMM contributed to management, processing and analyses of data. UL, KBW, LN, AMF, DB-F contributed to make data from the Lifebrain cohorts available. All authors contributed to reviewing and revising the manuscript draft.

