## Supplemental Information for "Brain aging differs with cognitive ability regardless of education"

**for Cognitive ability influences brain aging independently of education**

**Supplemental Methods**

*Supplemental Sample Information*

*Sample Recruitment, Inclusion and Exclusion Criteria, Demographics and Measures*

For all Lifebrain sub-cohorts (see detailed descriptions below), participants were only included if they had available MRI scans from at least two timepoints, and GCA score at minimum of one timepoint. For the UKB cohort, participants were only included if they had available MRI scans and GCA scores from at least two timepoints. While including also participants with only one time-point in analyses of level-level relationships of GCA and brain characteristics would yield more power, the current analysis was performed in the context of clarifying level-change relationships, and we thus wanted to reduce reliance on cross-sectional data not fully representative of longitudinal information(1).

*Lifebrain cohort*

Lifebrain cohorts BASE II, UB, Cam-CAN and Betula had MRI scans from two timepoints. For these, the same scanner was used at both timepoints. The LCBC cohort additionally had 38 participants with 3 scans, 95 with 4 scans, 37 with 5 scans, 1 with 6 scans, and 1 with 7 scans. Three different Siemens scanners were used for the Lifebrain cohort and due to the length of the follow-up interval, a portion (n = 148 of 411) were at some point followed up on a different scanner than the one used at baseline. For all but one of these participants, the scans were acquired at two scanners, whereas for one participant, 3 scanners were used in total. For most participants being followed up a different scanners, (n = 138), scans were acquired at both scanners used at the same timepoint (10 of whom at timepoint 2, 126 of whom at timepoint 3, and 2 of whom at timepoint 4). All scans were then included in analyses, and participants followed-up on different MRI scanners were independently processed for each scanner. Scanner was included as a covariate in all analyses. Demographics and number of participants from each Lifebrain sub-cohort are presented in Supplemental Table 1.

| **Study** | **Scanner** | **N** | **Time since baseline** | | | **Baseline age** | | | **Education** | | | **GCA at baseline** | | |
| --- | --- | --- | --- | --- | --- | --- | --- | --- | --- | --- | --- | --- | --- | --- |
|  |  |  | **M** | **SD** | **Range** | **M** | **SD** | **Range** | **M** | **SD** | **Range** | **M** | **SD** | **Range** |
| BASE II | Trio | 250 | 2.3 | 0.5 | 0.9-3.2 | 62.0 | 16.4 | 23.9-81.2 | 14.0 | 2.8 | 7-18 | 0.0 | 1.0 | -2.8-1.8 |
| LCBC | Avanto | 209 | 5.4 | 2.3 | 0.2-10.0 | 50.3 | 17.2 | 20.2-85.4 | 16.9 | 2.4 | 9-20 | 0.0 | 1.1 | -5.0-1.8 |
|  | Prisma | (4) | 3.6 | 0.4 | 3.1-4.0 | -- | -- | -- | -- | -- | --- | -- | -- | -- |
|  | Skyra | 202 | 4.6 | 3.6 | 0.2-11.0 | 48.3 | 22.5 | 20.0-81.9 | 16.1 | 2.6 | 9-20 | 0.0 | 0.9 | -3.9-1.6 |
| UB | Triotrim | 35 | 3.4 | 1.1 | 1.5-4.9 | 68.9 | 3.0 | 64.5-76.1 | 11.6 | 3.9 | 2-18 | 0.0 | 1.2 | -2.2-1.5 |
| Cam-CAN | Trio | 255 | 1.4 | 0.7 | 0.3-3.5 | 54.0 | 17.8 | 22.2-88.0 | 14.8 | 1.9 | 11-16 | 0.0 | 1.0 | -4.4-2.4 |
| Betula | Discovery | 178 | 4.6 | 0.5 | 4.0-5.0 | 59.9 | 13.4 | 25.5-80.8 | 13.4 | 3.7 | 6-20 | 0.0 | 1.7 | -2.8-2.5 |

***Supplemental Table 1. Overview of sample characteristics of included Lifebrain cohorts***

*Age, education, and time since baseline (scan) are given in years. GCA for Lifebrain is standardized per sample for first timepoint and given in z-scores. For LCBC, where different scanners were used, (N) refers to number of participants at the time the scanner was used, Prisma was not used at baseline.*

Below, we give further information on the origin of each cohort and the cohort-specific criteria and measures used.

**BASE II**

***Population, recruitment, inclusion/exclusion criteria and general description of study***

Participants of the Berlin Aging Study II (BASE II) were community-dwelling adults recruited from the greater Berlin metropolitan area through advertisements in newspapers and public areas (for cohort characteristics and additional details, see (2, 3)). Participants were invited to a medical exam consisting of a 2-day protocol, and two cognitive testing sessions scheduled 1 week apart, and were tested in small groups (e.g. about 6 participants per group) on a comprehensive cognitive battery that covers key cognitive abilities measured by 21 tasks. Each session lasted about 3.5 h. After completion of the cognitive examination of BASE-II, eligible participants were invited to take part in one MRI session within a time window of 2–4 weeks after cognitive testing. A subsample of the MR sample was later re-invited for follow-up. The different elements of the study were approved by the ethics committees of the Max Planck Institute for Human Development, the Charité University ethics committee and by the ethics committees of The German Association for Psychology (DGPs). Participants signed written informed consent and received monetary compensation for their participation in BASE-II and the MRI study. All experiments were performed in accordance with relevant guidelines and regulations.

***Inclusion/Exclusion Criteria/ Screening.*** An inclusion criterion for taking part in this study was age between 20 and 35 or 60 and 81 years at baseline entry, apparently healthy. Exclusion criteria were untreated diabetes and hypertension; prior stroke, head injuries or brain surgery; psychiatric illness; major depression; dementia with a score < 24 on the Mini-Mental State Examination. To that end, none of the participants took medication that might affect memory function or had a history of head injuries, medical (e.g., heart attack), neurological (e.g., epilepsy), or psychiatric disorders (e.g., depression). All participants reported normal or corrected to normal vision, were right-handed, and scored 27 or above on the Mini-Mental Status Examination.

***Variables Used for Education and General Cognitive Ability (GCA).*** Education was measured as number of years spent in formal schooling, also correcting for east/west German educational systems. The measures of fluid cognitive ability used for current GCA analyses, are the Practical Problems, Figural Analogies, and Letter Series tests, as described in (4).

**Betula**

***Population, Recruitment and General Description of Study/ Procedures.*** Betula was approved by the local ethics board at Umeå University. In the Betula longitudinal study on aging, memory and dementia, population-based sampling of healthy middle-aged and older adults was used for recruitment. Detailed recruitment procedures are found in (5, 6). For the current analyses, the MRI subsample of the study is used. Participation in the neuroimaging study was offered to all participants who had remained in the study and completed cognitive testing at the 5th Betula test wave in 2008-2009.

***Inclusion/Exclusion Criteria/ Screening.*** Exclusion criteria were severe visual or auditory handicaps, intellectual or developmental disabilities, suspected dementia, having a mother tongue other than Swedish, MRI contraindications, neurological disorders/conditions, or visual/motor deficits that could interfere with fMRI data collection, MMSE <24, brain or head surgery, and substantial brain anatomical deviations, scan movement artifacts, missing T1 images due to incomplete acquisition, or FreeSurfer processing failures.

***Variables Used for Education and General Cognitive Ability (GCA).*** Age at MRI-scanning, reported with a one decimal precision, was used in the analyses. Self-reported years of education with a half-year precision was used. Cognitive testing was performed 1-18 months prior to first scanning (mean interval: 9 months). For cognitive variables, fluid ability tests included the Block Design test (7), and a measure of immediate free recall of 16 enacted verb-noun sentences. Crystallized ability tests included a 30-item five-alternative forced choice vocabulary test, four measures of verbal fluency measured during one minute (generating words starting with the letter A, professions starting with the letter B, five-letter words starting with letter M, and words starting with F, without the letter A), as well as a 26-item general knowledge test. Exact testing procedures have been described in more detail elsewhere (5).

**Cam-CAN**

***Population, Recruitment and General Description of Study/ Procedures.*** Recruitment was done by invitation letters based on the patient lists of general practitioners within the Cambridge City area (for additional study information see(8) and <https://www.cam-can.org/>). A population-based cohort of 3000 adults aged 18 or above was recruited to Stage 1 of the project, where they completed an interview including health and lifestyle questions, a core cognitive assessment, and a self-completed questionnaire of lifetime experiences and physical activity. Of those interviewed, ~700 participants uniformly sampled across the lifespan aged 18-87 continued to Stage 2 where they undergo cognitive testing and provide measures of brain structure and function. A subset of adults returned for longitudinal follow-up data. The study is conducted in compliance with the Helsinki Declaration, and has been approved by the local ethics committee, Cambridgeshire 2 Research Ethics Committee (reference: 10/H0308/50).

***Inclusion/Exclusion Criteria/ Screening.*** General exclusion criteria: Term-time residents of colleges and universities, and participants whose Primary Care Physician feel are inappropriate to include. Exclusion criteria for the MRI part of the study: Not cognitively normal (MMSE < 24, memory defect, consent difficulties), communication difficulties (hearing problems [35db at 1000 Hz], insufficient English language, vision difficulties), medical problems by self-report of diagnosis (dementia diagnosis /Alzheimer’s Disease, Parkinson’s Disease, Motor Neuron disease, Multiple sclerosis, cancer, stroke, encephalitis, meningitis, epilepsy, head injury with serious results [coma, unconscious for >2 hours, skull fracture], recently diagnosed or uncontrolled high blood pressure, possible pregnancy, current psychiatric conditions [bipolar disorder, schizophrenia, psychosis]), mobility problems (restricted mobility which could prevent further participation, inability to walk 10 metres), substance abuse (past or current treatment for drug abuse, current drug usage), MRI/ MEG safety and comfort exclusions.

***Variables Used for Education and General Cognitive Ability (GCA).*** For fluid abilities, the standard form of the Cattell Culture Fair, Scale 2 Form A, was used(9). The Cattell test is a pen-and-paper test where the participant chooses a response on each trial from multiple choices, and records responses on an answer sheet. Our crystalized measure was the Spot The Word task (10). Spot the Word was time-limited to 5 minutes due to interview time restrictions but this was not indicated to the participants and interviewers could be flexible.

For education, we used catagorical values converted to years as follows: 1. College or university degree or higher (16), 2. A levels/AS levels or equivalent (13), 3. O levels/GCSEs or equivalent (11), 4. CSEs or equivalent (11), 5. NVQ or HND or HNC or equivalent (11), 6. Other professional qualifications e.g.: nursing, teaching (12), 0. None of the above (missing), 8. No answer (missing).

**LCBC**

***Population, Recruitment and General Description of Study/ Procedures*.** Cognitively healthy, community dwelling participants across the adult lifespan were drawn from studies of adults (age > 20 years) coordinated by the Research Group for Lifespan Changes in Brain and Cognition (LCBC [www.oslobrains.no](http://www.oslobrains.no)), approved by the Norwegian Regional Committee for Medical and Health Research Ethics South East. Written informed consent was obtained from all participants. The samples were recruited by newspaper and web page adds. Most participants were recruited for observational studies, while some adults were recruited to enter into cognitive training studies after baseline assessment.

***Inclusion/Exclusion Criteria/ Screening.*** Participants were screened using a standardized health interview prior to inclusion in the study. Participants with a history of self-reported neurological or psychiatric conditions, including clinically significant stroke, serious head injury, untreated hypertension, diabetes, and use of psychoactive drugs within the last two years, were excluded. Further, participants reporting worries concerning their cognitive status, including memory function, were excluded. All participants above 40 years scored ≥24 on the Mini Mental State Examination (11).

***Variables Used for Education and General Cognitive Ability (GCA).*** Education was recorded as total years of education to the highest obtained degree. For three persons with baseline age 20.3-21.3, only parental education was available, and was used in analyses. Age was recorded in years and months as the mean of age at the time of MRI scan and the age at the time of the cognitive testing. GCA was recorded based on the two-subtest (Vocabulary, Matrix reasoning) version of the WASI (12).

**UB**

***Population, Recruitment and General Description of Study/ Procedures*.** Recruitment and selection of participants *The WAHA Cohort (13)* from the University of Barcelona (UB) site took place between May 2012 and May 2014; the trial ended May 31, 2016. Eligible participants were recruited via mailing study brochures (LLU) or through the non-profit organization Institute of Aging (BCN), advertisements in the study centers, and word of mouth. Interested individuals attended an informational group meeting, completed a short medical questionnaire and signed the informed consent. Next candidates had a face-to-face interview with the study clinician, who assessed potential compliance, reviewed the medical history, inclusion and exclusion criteria, and recent blood work and use of medications or supplements, and administered the MMSE. Eligible participants were scheduled to have baseline tests (neuropsychological and ophthalmologic evaluations and collection of fasting blood and urine) and were then randomized to either a control or walnut group using a computerized random number table with stratification by center, sex, and age range. Couples entering the study were treated as one number and were randomized into the same group. The Bioethics Commission of the University of Barcelona (Institutional Review Board: IRB 00003099) approved the project.

***Inclusion/Exclusion Criteria/ Screening.*** Participants were healthy elderly men and women with normal cognitive and visual function. Inclusion criteria were age between 63 and 79 years, apparently healthy, and equally willing to be in either of the two groups (see below). Exclusion criteria included inability to undergo neuropsychological testing; morbid obesity (BMI ≥ 40 kg/m^2^); uncontrolled diabetes (HbA1c > 8%); uncontrolled hypertension (on-treatment blood pressure ≥ 150/100 mmHg); prior stroke, significant head trauma or brain surgery; relevant psychiatric illness; major depression; cognitive deterioration or dementia with a score < 24 on the Mini-Mental State Examination; other neurodegenerative disorders like Parkinson’s disease; advanced AMD or eye-related conditions precluding ophthalmological evaluation; prior chemotherapy; chronic illness with projected shortened lifespan; allergy to walnuts; customary use of fish oil and/or tree nuts (> 2 servings/week) and/or other relevant sources of ALA, such as flaxseed oil or soy lecithin.

***Variables Used for Education and General Cognitive Ability (GCA).*** Years of completed education was used. Block design from WAIS-III (14) and National Adult Reading Test (NART) (15) were used for GCA.

**UKB**

***Population, Recruitment and General Description of Study/Procedures.*** UK Biobank (UKB) (<https://www.ukbiobank.ac.uk/about-biobank-uk/>) is a major national and international health resource with the aim of improving the prevention, diagnosis and treatment of a wide range of illnesses. UK Biobank recruited ≈500,000 people aged between 40-69 years in 2006-2010 from across the country to take part in this project (16). Potential participants were identified through National Health Service (NHS) registers according to being aged 40-69 and living within a reasonable travelling distance of an assessment centre. Assessment centres (22 in total) are located in accessible and convenient locations with a large surrounding population. Participants have undergone measures and provided samples and detailed information about themselves and agreed to have their health followed. The dataset released in February 2020 was used. Age was calculated from year and month of birth (day of month is missing and was set to 1 for all subjects) to date of assessment. Age was calculated at the timepoint for the first MRI. Behavior data were collected at the timepoint for the MRI data collection).

***Inclusion/Exclusion Criteria/Screening.*** None of the participants included had a record of dementia of any kind.

***Variables Used for Education and General Cognitive Ability (GCA).*** Education: For the Biobank participants` generation, the UK school system provided free universal compulsory education between the ages of 5 and 15 to 16 years. A conversion table was made from the education categories: 1: College or University degree, 2: A levels/AS levels or equivalent, 3: O levels/GCSEs or equivalent, 4: CSEs or equivalent, 5: NVQ or HND or HNC or equivalent, 6: Other professional qualifications e.g.: nursing, teaching, -7: None of the above, -3: Prefer Not to answer. This was converted to years of education by taking the average value of the Data-Field 845, ”Age at completed education” minus 5 years, for all the participants in each category. Unfortunately, Date-Field 845 was not collected for education category 1: University/College. By taking this approach, we derived the following conversion: 1: 16, 2: 13, 3: 11, 4: 11, 5: 11, 6: 12, -7: 10, -3: missing. Education category 1, where we have very limited data, we set to it 16 years. General cognitive ability was entered as the raw fluid intelligence score (UKB Data-field 20016). This is a simple unweighted sum of the number of correct answers given to the 13 fluid intelligence questions.

***MRI scanning and processing*** Imaging data were collected and processed by the UKB (https://www.ukbiobank.ac.uk) as described in(17). Imaging data were collected using 3.0 T Siemens Skyra (32-channel head coil). Anatomical T1-weighted magnetization-prepared rapid gradient echo (MPRAGE) images were obtained in the sagittal plane at 1mm isotropic resolution, and T2 weighted FLAIR images were acquired at 1.05x1x1mm resolution in the sagittal plane. Images were cross-sectionally processed by the UK Biobank using the FreeSurfer 6.0 software package, and subsequently processed with the longitudinal stream of FS by the LCBC team, using FS version 6.0.

**Genetic analyses**

We obtained the Genetic Ancestry Factor (GAFs) estimated by the principal component analysis from the UK Biobank. We used the top 10 GAFs as covariates in our models to capture unmodeled ancestral difference among participants. The UK biobank sample was genotyped by the UK Biobank Axiom array from Affymetrix or the UK BilEVE array. The analysis only included participants that showed a kinship coefficient 0 (provided by the UK biobank). In addition, participants that failed the UKBiLEVE genotyping, or are outliers of heterozygosity estimates, or had high missingness were excluded from this analysis. The quality controls on variants has been performed by UK biobank and we did not perform further quality controls (for further details please see <https://www.ukbiobank.ac.uk/scientists-3/genetic-data/>). We computed the GAF for the remaining data based on SNPs that have MAF > 0.05 and are nearly independent, i.e., r-squared <0.1, using the PLINK program. To be consistent with UKB analysis, we included the top 10 GAFs as covariates in our statistical analysis. In PGSs analysis, GCA was first revisualized by GAFs by linear regression models. Then, we used the residuals GCA from such models in the analysis where PGSs for GCA and education(18, 19) were covariates. Note that as these PGSs were calculated based on the broader UKB sample, we do not investigate the effects of the PGSs themselves, but solely used them as covariates.

**Specific information on image acquisition in the different samples**

T1 weighted structural scans were acquired at Siemens, Philips and GE scanners at the various sites. The sequence parameters for each cohort are given in Supplemental Table 2.

| **Sample** | **Scanner** | **Field strength (T)** | **Sequence parameters** |
| --- | --- | --- | --- |
| BASE-II | Tim Trio Siemens | 3.0 | TR: 2500 ms, TE: 4.77 ms, TI: 1100 ms, flip angle: 7°, slice thickness: 1.0 mm, FoV: 256×256 mm, 176 slices |
| Betula | Discovery GE | 3.0 | TR: 8.2 ms, TE: 3.2 ms, TI: 450 ms, flip angle: 12°, slice thickness: 1 mm, FOV: 250×250 mm, 176 slices |
| Cam-CAN | Tim Trio  Siemens | 3.0 | TR: 2250 ms, TE: 2.98 ms, TI: 900 ms, flip angle: 9°, slice thickness 1 mm, FOV: 256×240 mm, 192 slices |
| LCBC | Avanto Siemens | 1.5 | TR: 2400 ms, TE: 3.61 ms, TI: 1000 ms, flip angle: 8°, slice thickness: 1.2 mm, FoV: 240×240 m, 160 slices, iPat = 2 |
|  | Avanto Siemens | 1.5 | TR: 2400 ms, TE = 3.79 ms, TI = 1000 ms, flip angle = 8, slice thickness: 1.2 mm, FoV: 240 x 240 mm, 160 slices |
|  | Skyra Siemens | 3.0 | TR: 2300 ms, TE: 2.98 ms, TI: 850 ms, flip angle: 8°, slice thickness: 1 mm, FoV: 256×256 mm, 176 slices |
|  | Prisma Siemens | 3.0 | TR: 2400 ms, TE: 2.22 ms, TI: 1000 ms, flip angle: 8°, slice thickness: 0.8 mm, FoV: 240×256 mm, 208 slices, iPat = 2 |
| UB | Tim Trio Siemens | 3.0 | TR: 2300 ms, TE: 2.98, TI: 900 ms, slice thickness 1 mm, flip angle: 9°, FoV: 256×256 mm, 240 slices |
| UKB | Skyra  Siemens | 3.0 | TR: 2000 ms, TI: 880 ms, slice thickness: 1 mm, FoV: 208×256 mm, 256 slices, iPAT=2 |

**Supplemental Table 2** MR acquisition parameters

TR: Repetition time, TE: Echo time, TI: Inversion time, FoV: Field of View, iPat: in-plane acceleration, GRAPPA: GRAPPA acceleration factor. *Customized

**Statistical analyses**

In order to obtain PCA estimates of cognition from missing data, we had to impute missing data. To do so we used the iterativeImputer Class from the scikit-learn (v1.0.1) python package ([Scikit-learn: Machine Learning in Python](http://jmlr.csail.mit.edu/papers/v12/pedregosa11a.html))(20), which models each feature with missing values as a function of other features, and uses subsequent estimates for imputation.

Contact information for data requests:

| **Study** | **Lifebrain PI** | **e-mail** |
| --- | --- | --- |
| BASE-II | Ulman Lindenberger | |
| BETULA | Lars Nyberg | |
| Cam-CAN | Rik Henson | |
| LCBC | Kristine B Walhovd | |
| UB | David Bartres-Faz | |

**Supplemental Results**

**Associations of general cognitive ability and cortical characteristics**

**A. Cortical volume**

**UKB**

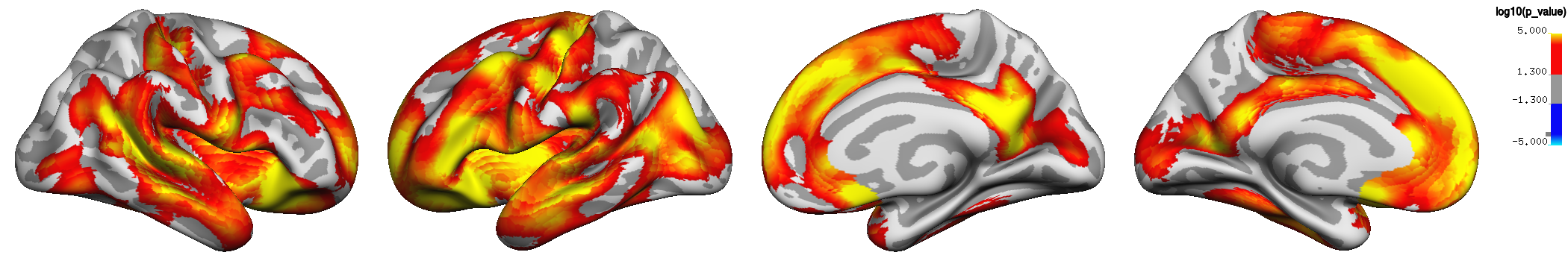

**Lifebrain**

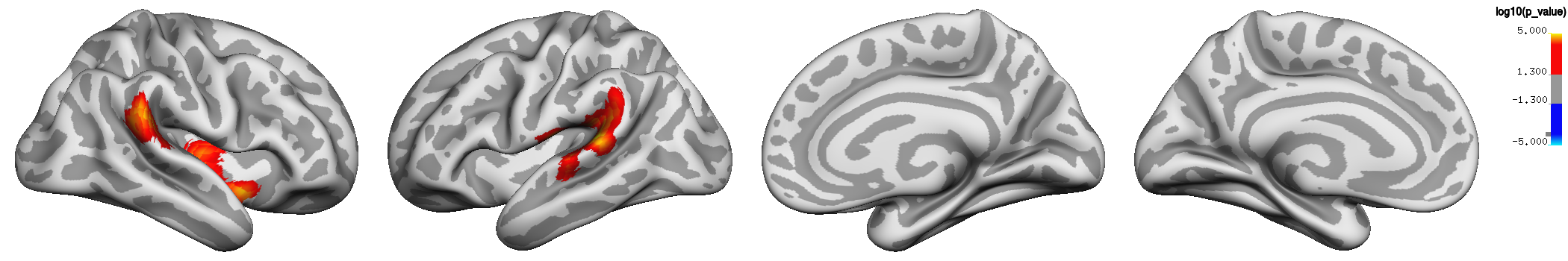

**B. Cortical area**

**UKB**

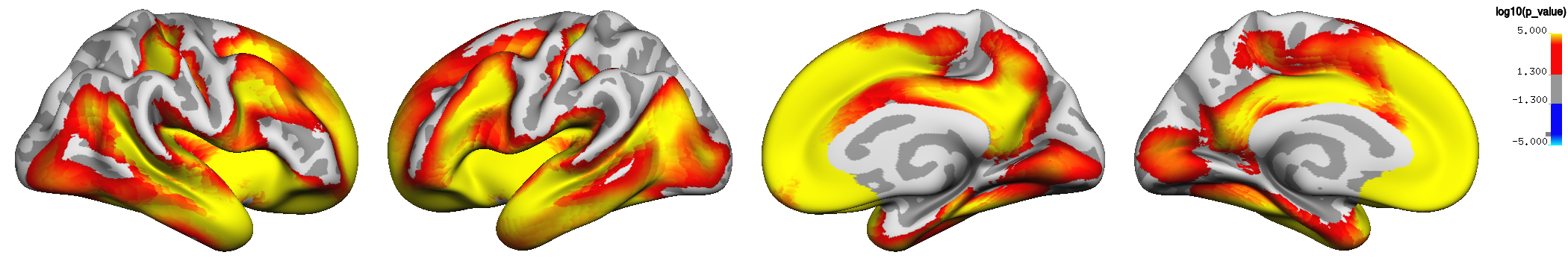

**Lifebrain**

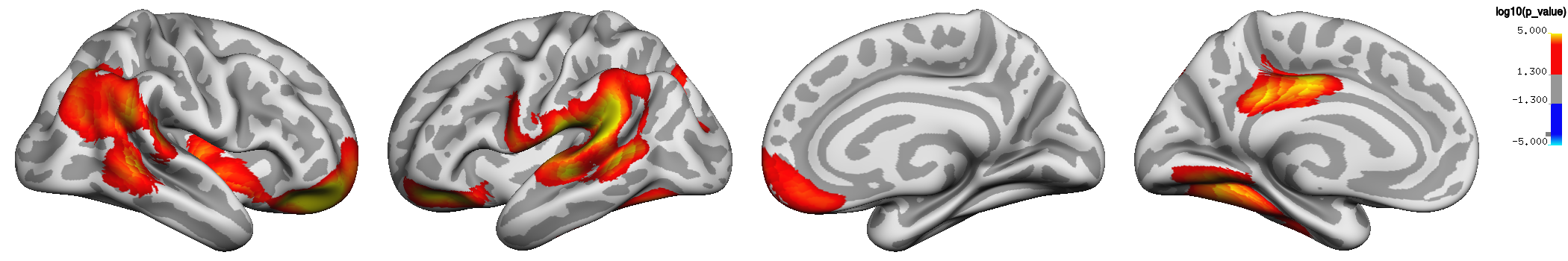

**C. Cortical thickness**

**UKB**

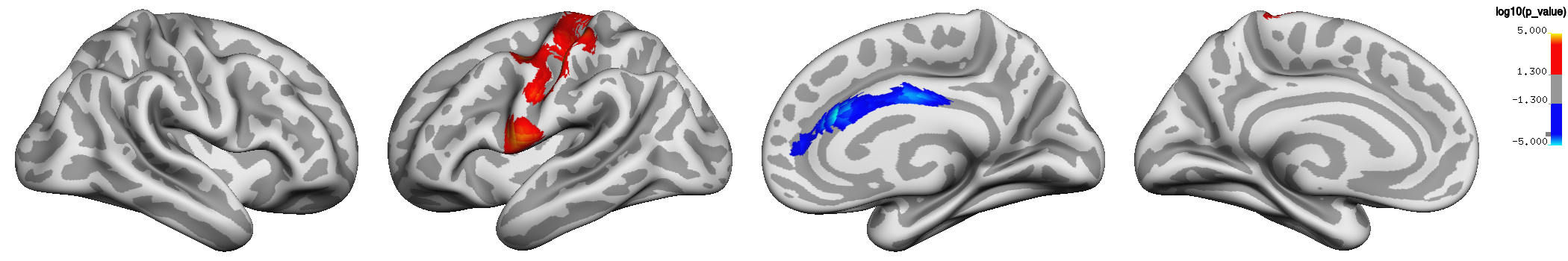

**Lifebrain**

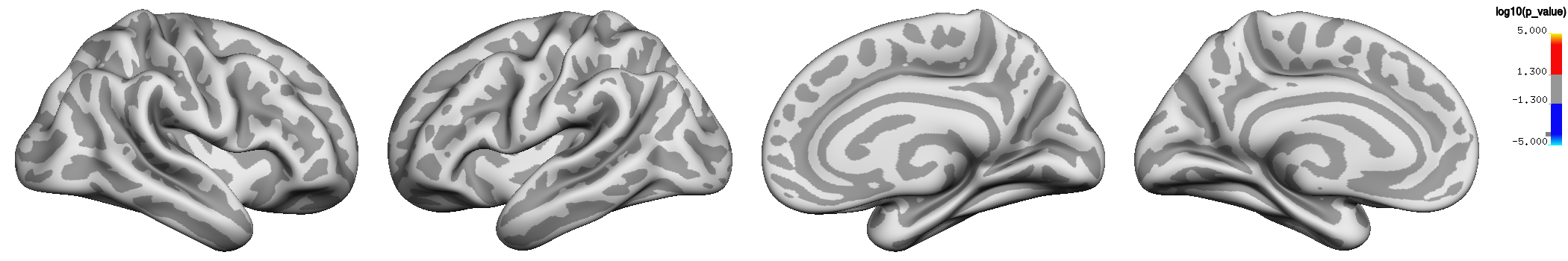

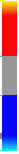
 Log10 (p-value)

-5.0 -1.3	1.3	5.0

***Supplemental Figure 1.*** *P-value maps of* ***the associations of general cognitive ability (GCA) and cortical characteristics when education is not controlled for.***

*The relationships between GCA at baseline and cortical characteristics are shown, when age, sex, and time (since first scan) are controlled for (p <.01, corrected using a cluster-forming p-value threshold of p < .01). Effects are shown, from left to right for each panel: right and left lateral view, right and left medial view, for UKB and Lifebrain samples for A. Cortical volume, B. Cortical area, and C. Cortical thickness.*

**Associations of general cognitive ability and change in cortical characteristics**

**controlled for the interaction of age by time**

**A. Cortical volume**

**UKB**

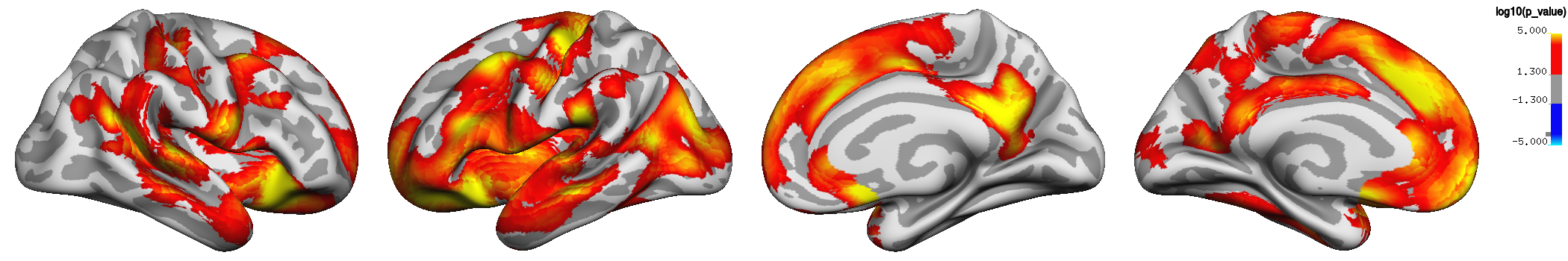

**Lifebrain**

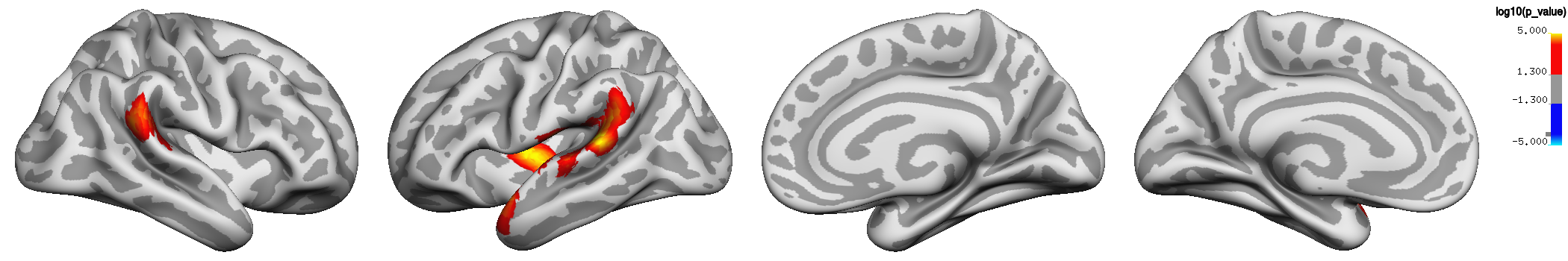

**B. Cortical area**

**UKB**

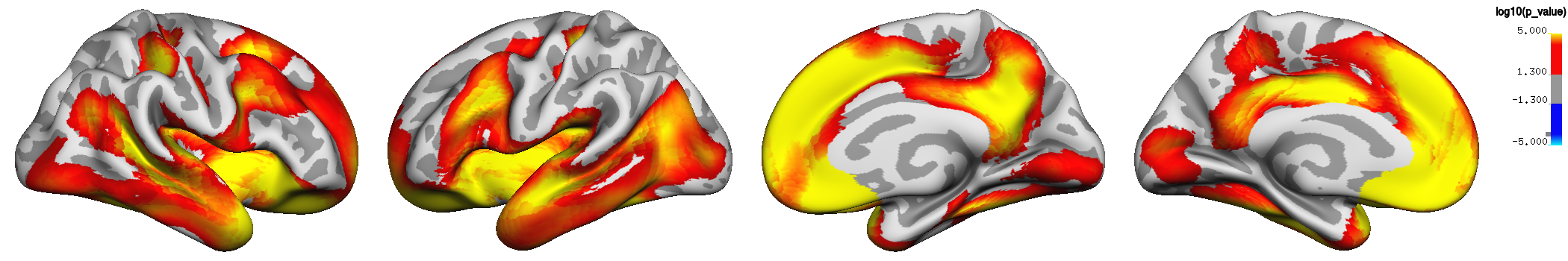

**Lifebrain**

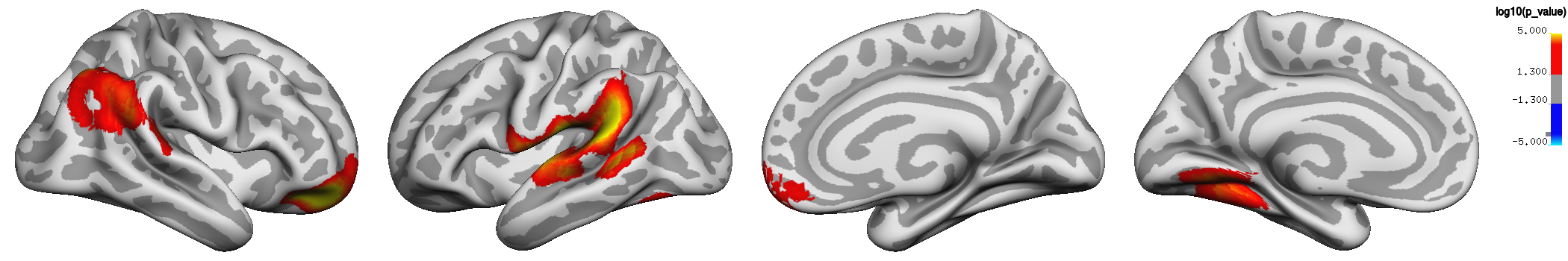

**C. Cortical thickness**

**UKB**

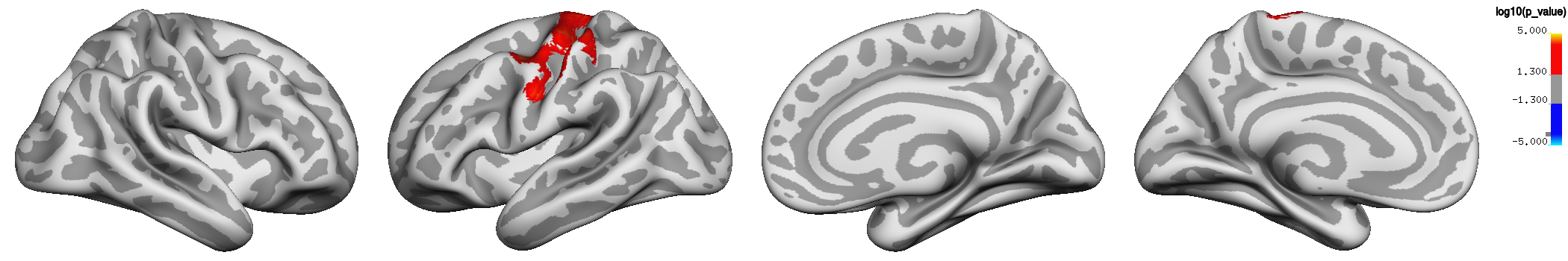

**Lifebrain**

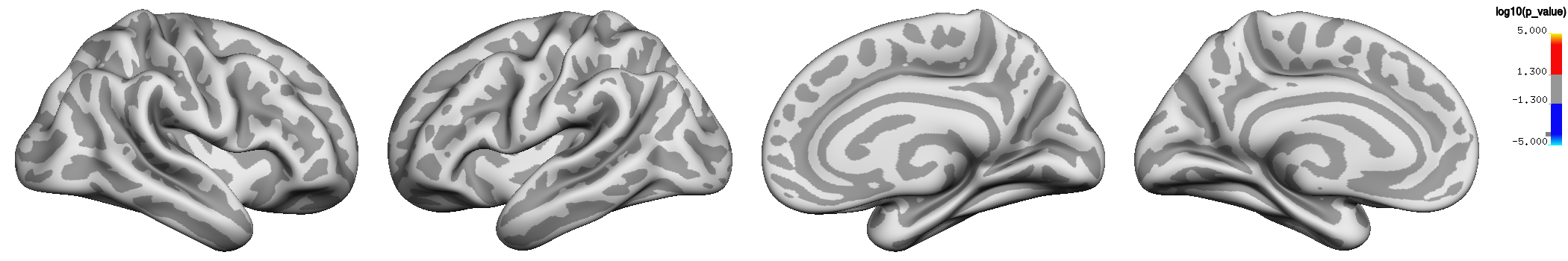

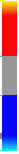
 Log10 (p-value)

-5.0 -1.3	1.3	5.0

**Supplemental Figure 2. P-value maps of the associations of general cognitive ability (GCA) and cortical characteristics, controlling for age interactions.** The associations are shown for GCA at baseline, when age, sex, education, time (since baseline scan), and the interaction of age at baseline and time, are controlled for (p <.01, corrected using a cluster-forming threshold of p < .01). Significant regions are shown, from left to right for each panel: right and left lateral view, right and left medial view, for UKB and Lifebrain samples for A. Cortical volume, B. Cortical area, and C. Cortical thickness.

**Associations of general cognitive ability and cortical characteristics**

**controlled for the effect of education *and ICV* in the Lifebrain sample**

**A. Cortical volume**

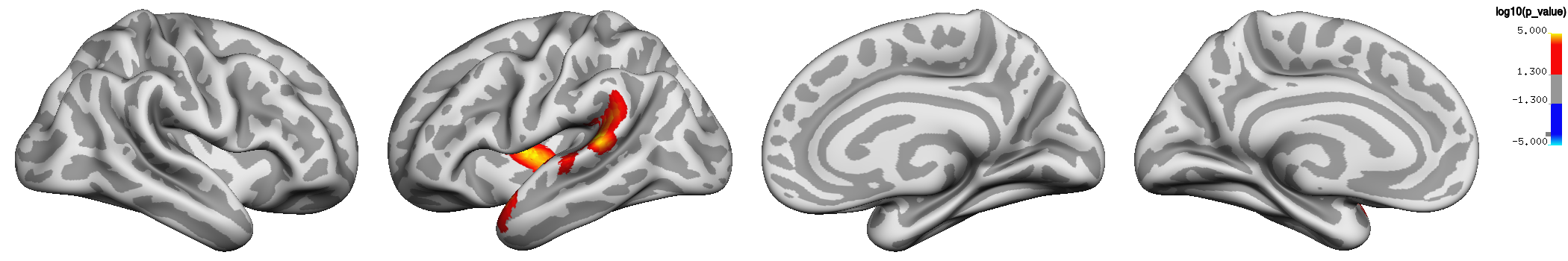

**B. Cortical area**

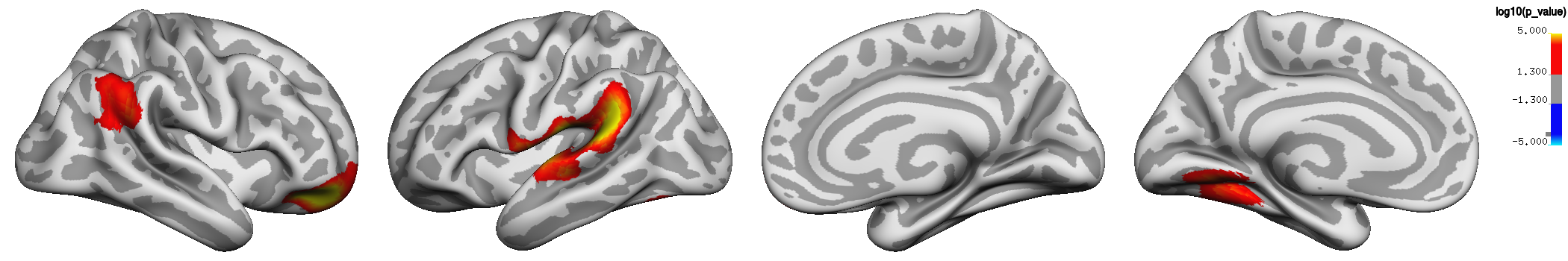

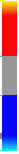
 Log10 (p-value)

-5.0 -1.3	1.3	5.0

***Supplemental Figure 3. P-value maps of the associations of general cognitive ability (GCA) and cortical characteristics, controlled for education and Intracranial Volume (ICV).***

*Associations of GCA at baseline are shown, when age at baseline, sex, time (since baseline scan), education and ICV are controlled for (p <.01, corrected using a cluster-forming p-value threshold of p<.01). Significant regions are shown, from left to right for each panel: right and left lateral view, right and left medial view, for Lifebrain samples for A. Cortical volume, and B. Cortical area. No effects were seen for cortical thickness, and no effects were seen in UKB.*

**Associations of general cognitive ability and change in cortical characteristics**

**controlled for the effect of education, p <.05**

A. **Cortical volume**

**UKB**

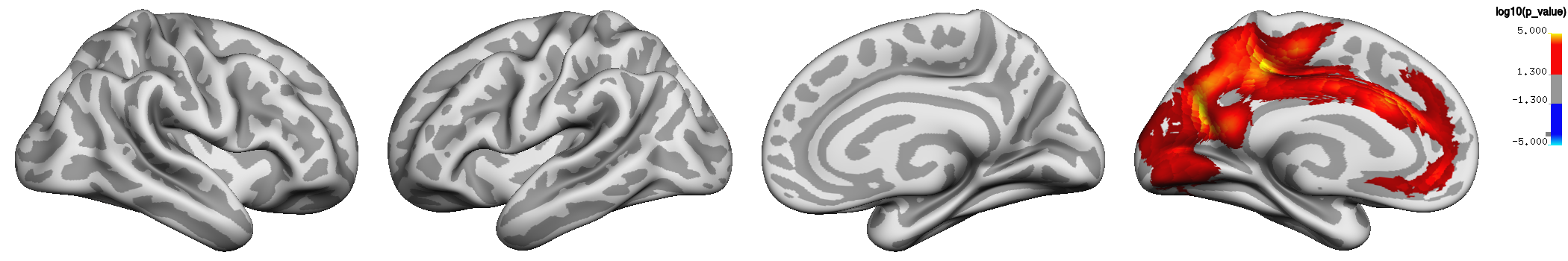

**Lifebrain**

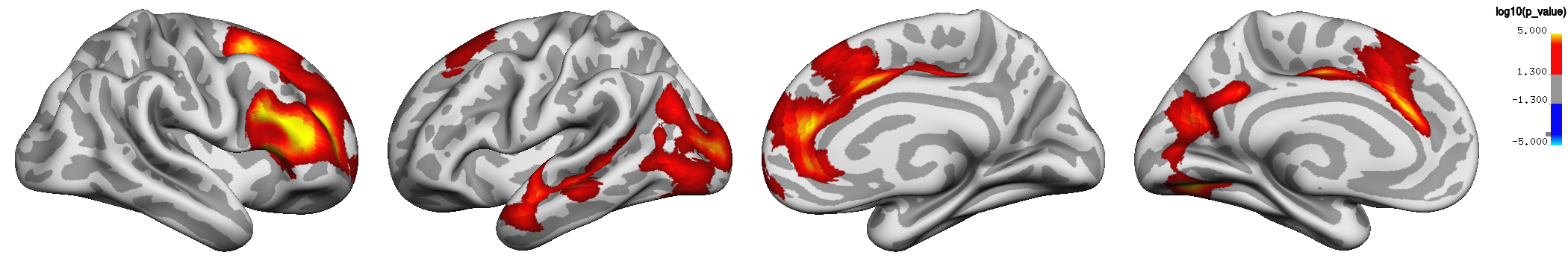

B. **Cortical area**

**UKB**

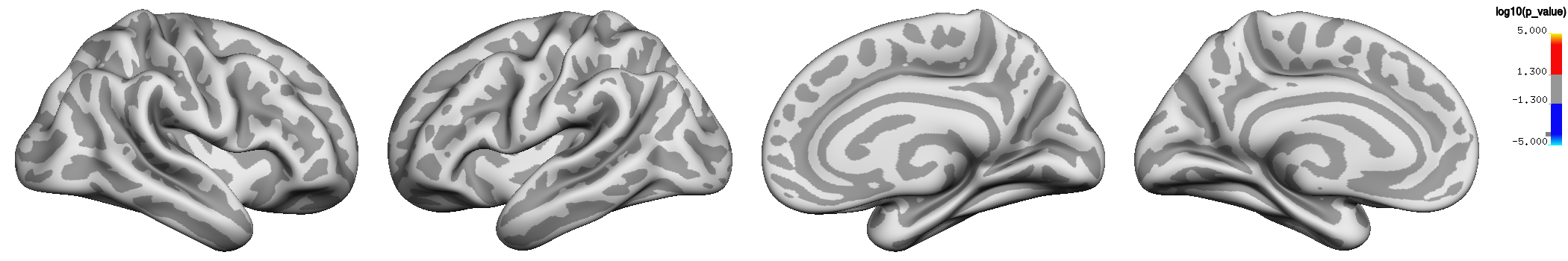

**Lifebrain**

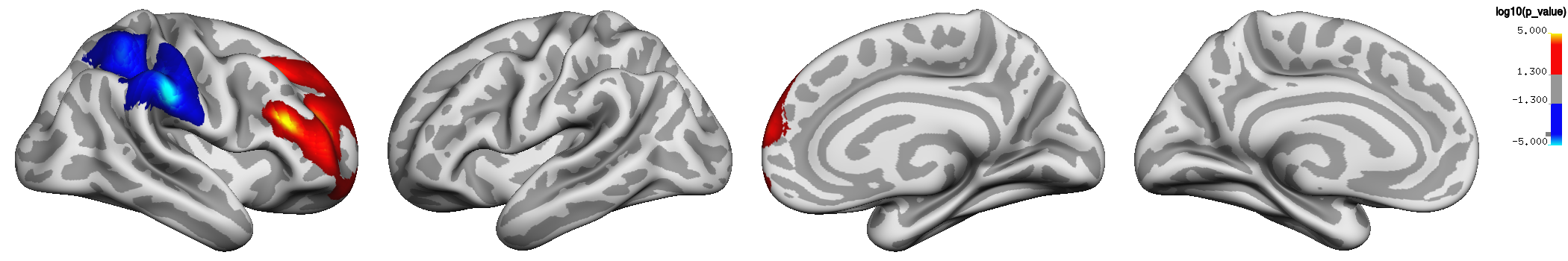

**C. Cortical thickness**

**UKB**

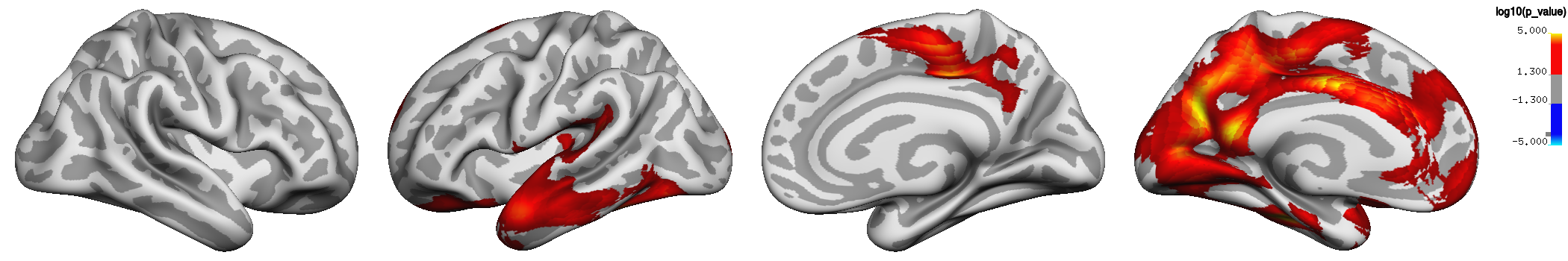

**Lifebrain**

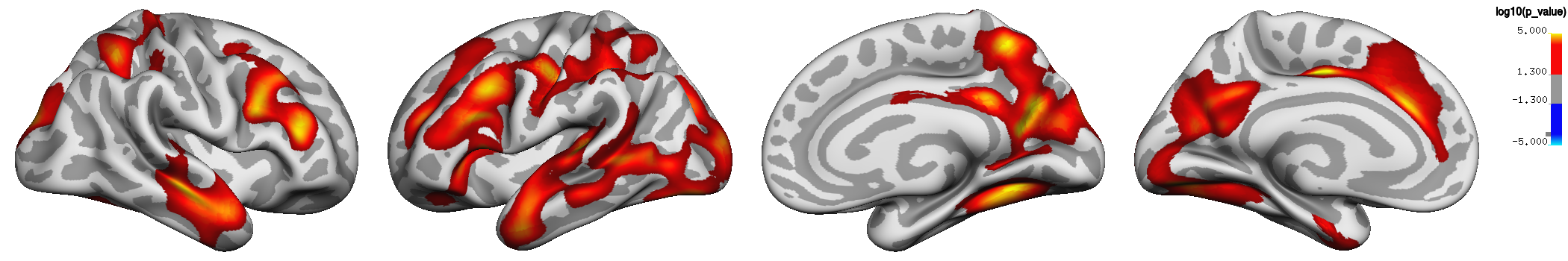

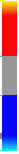
 Log10 (p-value)

-5.0 -1.3	1.3	5.0

**Supplemental Figure 4. P-value at p <.05 (corrected using a cluster-forming p-value threshold of p<.05) for the associations of general cognitive ability (GCA) and change in cortical characteristics, controlling for the effect of education over time.**

The associations are shown for the interaction of GCA at baseline and time (interval since baseline scan), when age, sex, time, GCA, education, and the interaction of education by time, are controlled for. Compare to Figure 2 (showing results at p-value p <.01, corrected using a cluster-forming p-value threshold of p<.01). Significant regions are shown, from left to right for each panel: right and left lateral view, right and left medial view, for UKB and Lifebrain samples for A. Cortical volume, B. Cortical area, C. Cortical thickness.

**Associations of general cognitive ability and change in cortical characteristics**

**controlled for the effect of education and ICV**

A. **Cortical volume**

**UKB**

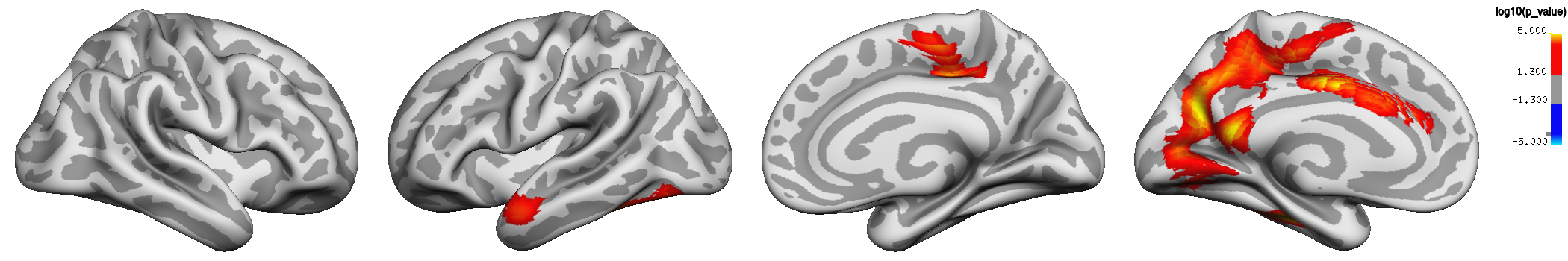

**Lifebrain**

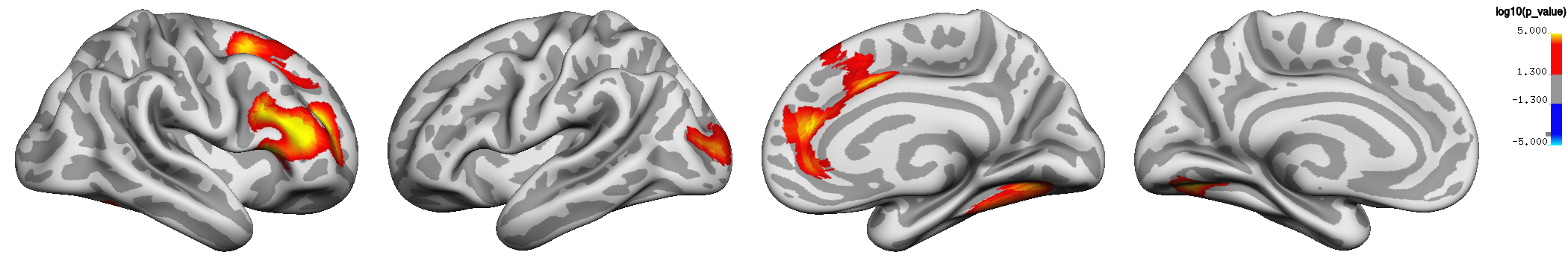

**B. Cortical area**

**UKB**

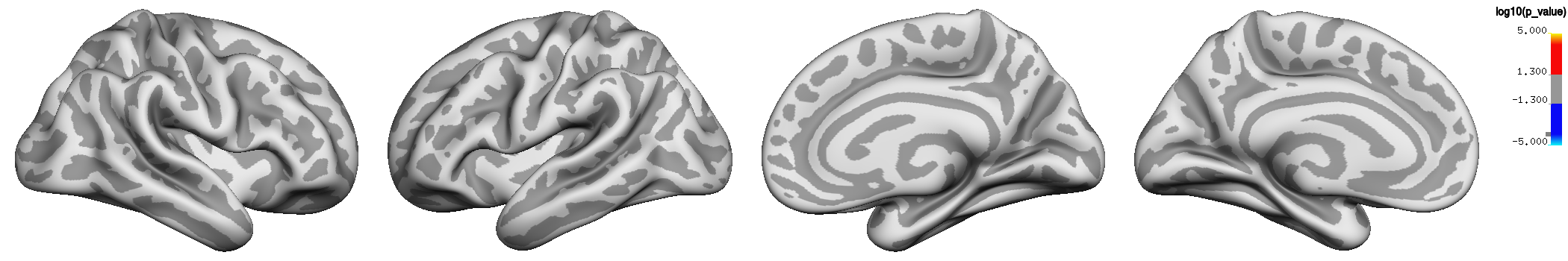

**Lifebrain**

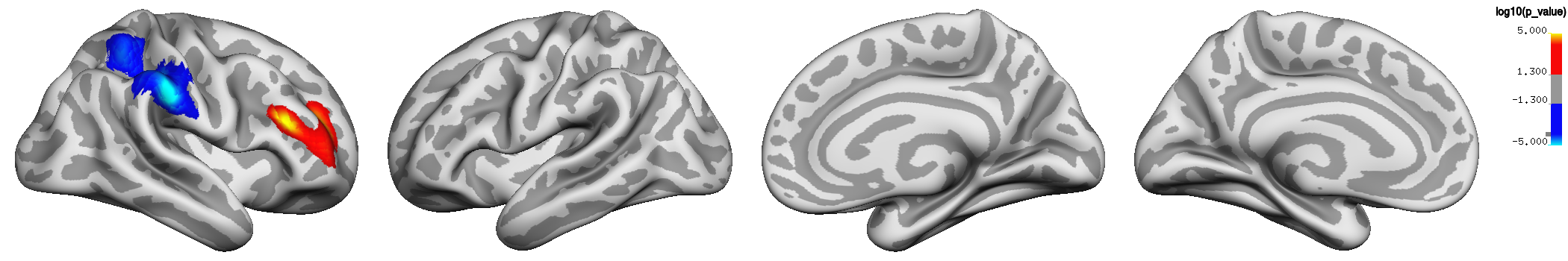

**C. Cortical thickness**

**UKB**

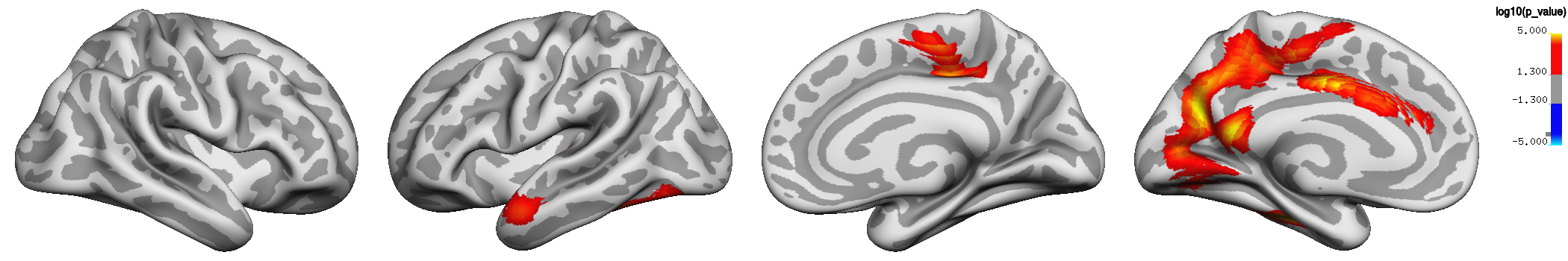

**Lifebrain**

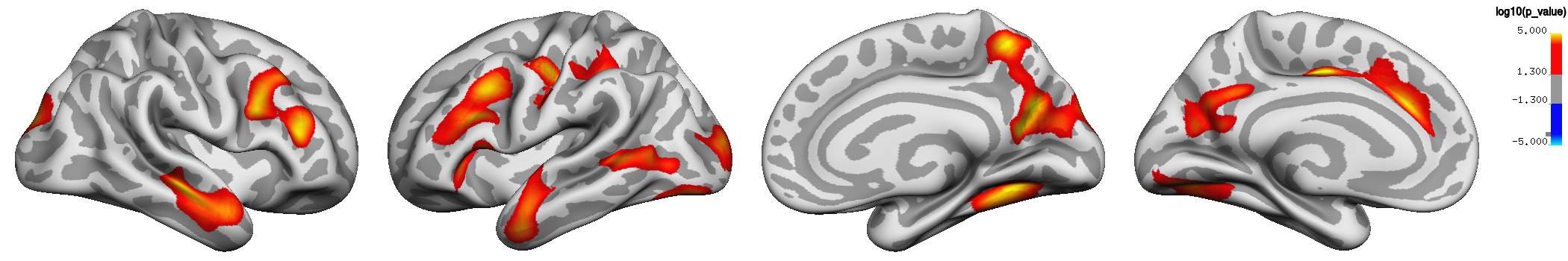

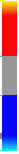
 Log10 (p-value)

-5.0 -1.3	1.3	5.0

***Supplemental Figure 5. P-value maps of associations of general cognitive ability (GCA) and change in cortical characteristics, controlled for the effect of education over time, and for intracranial volume (ICV).*** *The associations are shown for the interaction of GCA at baseline and time (interval since baseline scan), when age, sex, time, GCA, education, the interaction of education and time, and ICV, are controlled for (p <.01, corrected using a cluster-forming p-value threshold of p<.01). Effects are shown, from left to right for each panel: right and left lateral view, right and left medial view, for UKB and Lifebrain samples: A. Cortical volume, B. Cortical area, C. Cortical thickness. The inclusion of ICV in these analyses essentially did not affect the results.*

**Associations of general cognitive ability (GCA) and cortical characteristics**

**controlled for education and polygenic scores (PGSs) for education and GCA**

A. **Cortical volume**

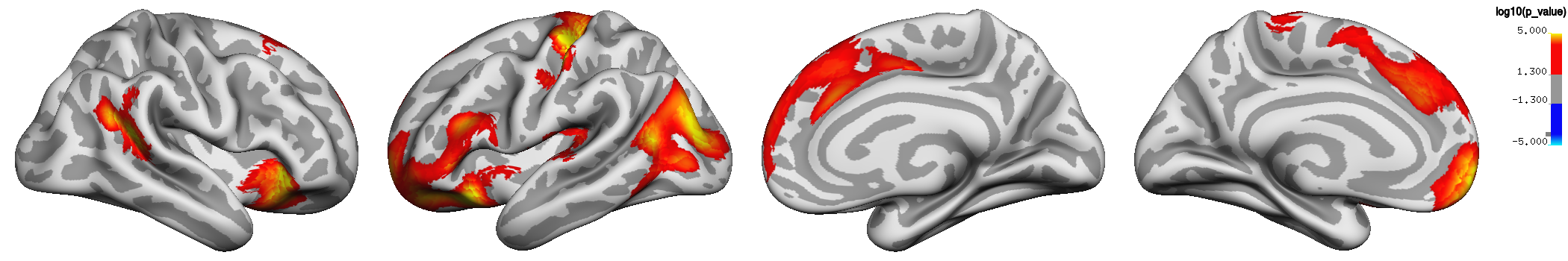

B. **Cortical area**

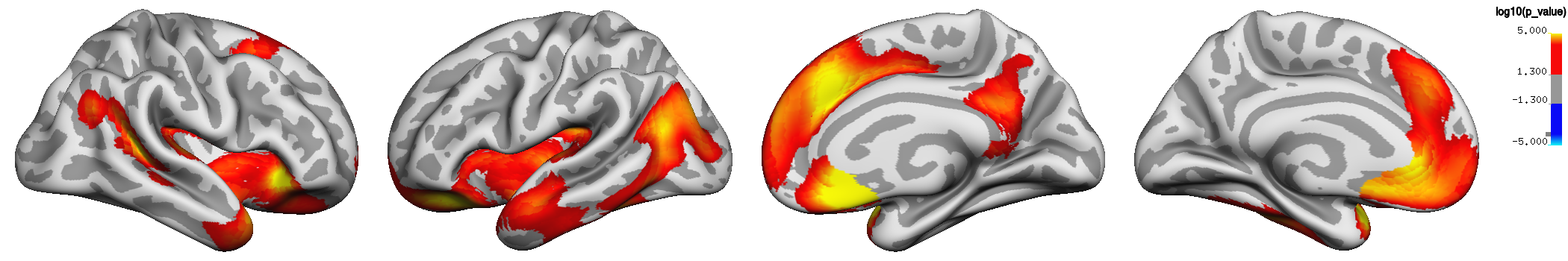

**C. Cortical thickness**

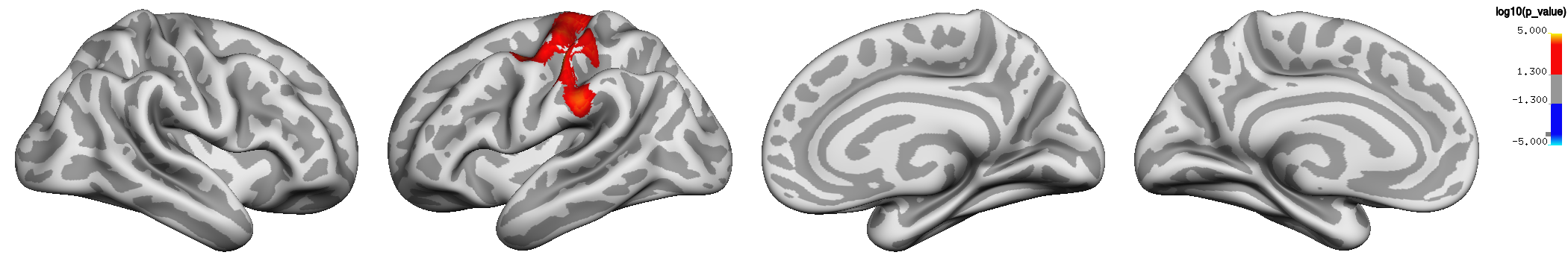

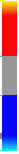
 Log10 (p-value)

-5.0 -1.3	1.3	5.0

***Supplemental Figure 6. P-value maps of associations in the UKB of general cognitive ability (GCA) and cortical characteristics, controlled for education and polygenic scores (PGSs) for education and GCA***

*Associations of GCA at baseline are shown, when age at baseline, sex, time (interval since baseline scan), education and PGSs for education and GCA are controlled for (p <.01, corrected using a cluster-forming p-value threshold of p<.01). Significant regions are shown, from left to right for each panel: right and left lateral view, right and left medial view, for the UKB and the LCBC samples for A. Cortical volume, B. Cortical area, and C. Cortical thickness.*
